# Integrative Multi-omic Profiling of Two Human Decedents Receiving Pig Heart Xenografts Reveals Strong Perturbations in Early Immune-Cell and Cellular Metabolism Responses

**DOI:** 10.1101/2023.06.05.543406

**Authors:** Eloi Schmauch, Brian Piening, Bo Xia, Chenchen Zhu, Jeffrey Stern, Weimin Zhang, Alexa Dowdell, Bao-Li Loza, Maede Mohebnasab, Loren Gragert, Karen Khalil, Brendan Camellato, Michelli Faria de Oliveira, Darragh O’Brien, Elaina Weldon, Xiangping Lin, Hui Gao, Larisa Kagermazova, Jacqueline Kim, Alexandre Loupy, Adriana Heguy, Sarah Taylor, Florrie Zhu, Sarah Gao, Divya Gandla, Kriyana Reddy, Andrew Chang, Basil Michael, Lihua Jiang, Ruiqi Jian, Navneet Narula, Suvi Linna-Kuosmanen, Minna Kaikkonen-Määttä, Marc Lorber, Manolis Kellis, Vasishta Tatapudi, David Ayares, Adam Griesemer, Massimo Mangiola, Harvey Pass, Michael P. Snyder, Robert A. Montgomery, Jef D. Boeke, Brendan J. Keating

## Abstract

**Background:** Recent advances in xenotransplantation in living and decedent humans using pig xenografts have laid promising groundwork towards future emergency use and first in human trials. Major obstacles remain though, including a lack of knowledge of the genetic incompatibilities between pig donors and human recipients which may led to harmful immune responses against the xenograft or dysregulation of normal physiology. In 2022 two pig heart xenografts were transplanted into two brain-dead human decedents with a minimized immunosuppression regime, primarily to evaluate onset of hyper-acute antibody mediated rejection and sustained xenograft function over 3 days.

**Methods:** We performed multi-omic profiling to assess the dynamic interactions between the pig and human genomes in the first two pig heart-xenografts transplants into human decedents. To assess global and specific biological changes that may correlate with immune-related outcomes and xenograft function, we generated transcriptomic, lipidomic, proteomic and metabolomics datasets, across blood and tissue samples collected every 6 hours over the 3-day procedures.

**Results:** Single-cell datasets in the 3-day pig xenograft-decedent models show dynamic immune activation processes. We observe specific scRNA-seq, snRNA-seq and geospatial transcriptomic changes of early immune-activation leading to pronounced downstream T-cell activity and hallmarks of early antibody mediated rejection (AbMR) and/or ischemia reperfusion injury (IRI) in the first xenograft recipient. Using longitudinal multiomic integrative analyses from blood in addition to antigen presentation pathway enrichment, we also observe in the first xeno-heart recipient significant cellular metabolism and liver damage pathway changes that correlate with profound physiological dysfunction whereas, these signals are not present in the other xenograft recipient.

**Conclusions:** Single-cell and multiomics approaches reveal fundamental insights into early molecular immune responses indicative of IRI and/or early AbMR in the first human decedent, which was not evident in the conventional histological evaluations.

## INTRODUCTION

Organ shortages remain a major limitation within human allo-transplantation, with waiting lists greatly outpacing the number of donors available. Genetically modified pig organs offer several significant advantages for transplantation into humans and may help alleviate the current critical shortage of suitable organs ^1,2^. Advances in ethical research using recently deceased brain-dead human donors, and genetic knockout models of key xeno-antigens including *alpha1,3-galactosyltransferase (α-1,3-Gal)* has enabled the first sets of pig to non-human primate xenotransplants to be performed ^3,4^.

On September 25 and November 22 2021, pig thymus-kidney (“thymokidney”) xenografts were transplanted to two decedents ^5^ and on September 30^th^, 2021 an independent group transplanted two pig kidneys into a human decedent ^6^. In June and July 2022 two pig heart xenografts were transplanted into two human decedents with the primary aims being to assess the presence of hyper-acute antibody mediated rejection and sustained xenograft functioning over a 3-day protocol. The immunosuppression regimen did not include costimulatory blockade drugs which are considered essential to support long term survival of a xenograft ^7,8^. The 10-gene-edit pig heart xenografts received by both decedents showed no ostensible evidence of cellular or antibody-mediated rejection on conventional histology, or on flow or complement-dependent cytotoxicity crossmatches (Moazami *et al*. in press). Data on systemic responses, hemodynamic stability, and cardiac performance were collected over 66 hours post-transplant. Decedent 1 received a heart that was undersized for the recipient and had a longer cold ischemic time in static storage than decedent 2. The hemodynamics and left ventricular stroke volume of decedent 1 began to deteriorate at 30-36 hours after reperfusion, whereas the cardiac function of decedent 2 remained stable throughout the study. Decedent 1 developed an increased need for vasopressors which lead to evidence of visceral hypoperfusion including a rising lactate and liver enzymes indicating ischemic injury to end organs. The explant tissue from decedent 1 demonstrated progressive myocyte injury and cell death not present in day 1 and 2 endomyocardial biopsies. This was not observed in the xenograft of decedent 2. Standard histology and immunohistology did not show features of hyperacute rejection or antibody mediated rejection in either explant or daily biopsies.

Longitudinal high-dimensional multiomics profiling approaches developed over the last decade have been powerful tools for identifying novel biomolecular signatures across a range of biological processes and disease states, including obesity, stress, infection, pregnancy, post-transplant outcomes, metabolic disease and in extreme physiological states such as prolonged periods of human space flight ^9-15^. One of the key benefits of the longitudinal nature of this approach is that time points from each individual can serve as their own internal controls, resulting in orders-of-magnitude gains in statistical power vs standard cross-sectional studies^11^. Moreover, the recent development of single cell geo-spatial transcriptomics studies enables accurate assessment of mRNA expression in a given cellular state and cellular composition changes (single cell RNA-seq), in addition to their spatial co-localization and cross-talk (geospatial RNA-seq) ^16,17,18^.

We performed longitudinal multi-omic profiling to assess the dynamic interactions between the pig and human genomes in the first two pig heart-xenografts transplants into human decedents. In attempts to gain biological insights into these key xenotransplant procedures, we generated lipidomic, proteomic and metabolomics as well as bulk-RNA-seq and single-cell/nuclei RNA-seq and geospatial transcriptomic datasets across blood and tissue samples collected throughout the 3-day xenotransplant protocols. To assess global changes, we performed system-level analysis through integration of the respective -omics datasets.

## MATERIALS AND METHODS

### Patient Data and Biospecimens collections

The two human decedents recipients of the pig heart xenografts were one male and one female, both of European ancestry. Consent and other regulatory protocols, operative procedures, available phenotypes and physiological and clinical outcomes are described in detail elsewhere (Moazami *et al*. in press Nature Medicine). Key biospecimens as well as physiological measurements, blood chemistries and other lab values are described in **Supplementary Table 1**. Blood samples were collected at the following timepoints for decedent 1: pre-transplant (0 h) and then at 6, 12, 18, 24, 30, 36, 42, 48, 54, 60, 65, 66 and 66.5 h (terminal sample) and for decedent 2 at: 0,6, 12, 18, 24, 30, 36, 42, 48, 54, 60 and 66 h (terminal sample).

### Transcriptomics

RNA profiling was performed using different biospecimen sources across the two pig- to human decedent cardiac xenotransplant procedures, spanning PBMC and fresh-frozen tissue samples using multiple methodologies/processes including bulk RNA-seq, targeted RNA profiling, single-cell (sc)- and single-nuclei(sn)-RNA-seq and geospatial transcriptomics as described below. Bulk- and scRNA-seq was performed on blood samples collected at the timepoints indicated above except for the last three bulk RNAseq timepoints of decedent 1 which were collected at 60, 64 and 65 hrs.

<u>Bulk RNA-seq</u> was performed primarily to assess the concordance with the scRNA-seq as the latter methodologies is more prone to batch effect with a smaller number of starting input cells leading to wider variance in cell types. These analyses were performed using venous blood collected into PaxGene tubes from the decedent and processed as a single batch (all timepoints across both decedents). For data analysis, FASTQ files were aligned to GRCh38 GENCODE Human release v43 using *STAR* v2.7-10a. *STAR* run parameters were adapted from the ENCODE RNA-Seq pipeline for gene count quantification. The counts were preprocessed by removing genes with less than an average of 4 counts across all samples. Differential gene expression analysis was performed on each contrast using *DESeq2*.

<u>For scRNA-seq of PBMCs</u> we used a standard 10x Genomics scRNA-seq protocol (10x Genomics, CA, USA). Initial scRNA-seq data processing was performed using the *CellRanger* pipeline using the vendor-recommended workflow. After *cell-ranger* preprocessing, the count matrices were processed using *scanpy*^19^. In brief, QC filters were applied as follows: maximum 50,000 counts; maximum 7,000 genes; maximum 20% mitochondrial gene expression; minimum 500 genes. Several levels of corrections were applied: regression on the number of unique molecular identifiers (UMIs), genes and cell cycle. In addition, *Harmony*^20^ was used for batch correction, at the subject level (to integrate the two decedents) for main cell-type embedding, and at the timepoint level to correct for technical batch-effect for subtypes analysis. The counts were normalized and logarithmized. For visualization and clustering, highly variable genes are selected using *scanpy’s* dedicated function, and the integrated matrix was scaled. Then PCA and UMAP algorithm was applied (including the UMAP *knn graph creation*). From that knn graph, clusters are discovered using the leiden algorithm. Cell-types are discovered based on marker genes and gene set enrichment.

<u>For snRNA-seq</u> the starting material was 4 × 50μm OCT curls, from which nuclei were isolated using Chromium Nuclei Isolation Kit (https://cdn.10xgenomics.com/image/upload/v1660261285/support-documents/CG000505_Chromium_Nuclei_Isolation_Kit_UG_RevA.pdf). The isolated nuclei were resuspended in 50μl of wash and resuspension buffer, counted using an automated cell counter, and immediately loaded into Gel Bead-in-Emulsion (GEMs) as a single replicate and run according to the Chromium Single Cell 3’ kit (https://cdn.10xgenomics.com/image/upload/v1668017706/support-documents/CG000315_ChromiumNext GEMSingleCell3-_GeneExpression_v3.1_DualIndex RevE.pdf). For geospatial transcriptomics, we used 10μm OCT sections in duplicate (serial sections processed to single nuclei curls). Fixation, hematoxylin and eosin (H&E) staining and imaging for the Visium processing was performed using the Visium v1 protocol (CG000239) (https://cdn.10xgenomics.com/image/upload/v1660261285/support-documents/CG000160_DemonstratedProtocol_MethanolFixationandHEStaining_RevC.pdf). For both single-nuclei RNA-seq (snRNA-seq) and geospatial transcriptomics (Visium, 10x Genomics) we first embedded a flash frozen samples in Optimal Cutting Temperature (OCT) compound to preserves the structure of the pig heart xenograft tissue and to provide support during cryo-sectioning, and they were kept at -80°C prior to sectioning.

For snRNA-seq and geospatial, the raw data was processed using *Cellranger* with mixed genomes (‘Barnyard’ experiment). This enabled the correct labelling and assignment of pig and human nuclei in addition to cross-species multiplet removal (**Supplementary Figure 1**). To ensure the absence of multiple alignments, *Cellranger* was additionally run against the human genome (GRCh38) and pig genome (Sscrofa11) separately. For further analysis, the data were partitioned by species. Due to the low number of human cells, lenient filtering was applied (100 minimum genes, 200 minimum counts, 20% maximum mitochondrial content). For the pig nuclei, we filtered with the following parameters: the maximum number of UMI was 15,000, the minimum number of genes was set to 500, while the maximum set to 4,000, the maximum mitochondrial content was set to 5%, and the minimum number of counts deemed acceptable was 500). Counts were normalized, logarithmized, highly variable genes selected, with the data corrected through regression by number of UMI and mitochondrial content. The same approach then the PBMC scRNA-seq was then utilized (*PCA, UMAP, Leiden*).

### Proteomics

Proteomic profiling for all time points of the plasma samples was performed by 16-Plex Tandem Mass Tag (TMT) based Mass Spectrometry (MS). A pool of all samples was used as a reference channel across different runs. Protein quantification and QA/QC were performed as previously described^19^. Briefly, in each MS run, Peptide Spectrum Matches (PSMs) without quantification were removed before sum normalization with the reference channel. Only PSMs with unique protein groups were selected which had a SPS match of >65, and a co-isolation interference of <50. PSMs with a sum of 200 from all 16 channels were removed. The resulting PSMs were combined into protein groups. Blanks values were imputed with the median of the smallest quantity in each run. All four runs were then combined, and the ratios of samples to reference channel were calculated. In total, over 1000 protein groups were identified (**Supplementary Figure 2**).

### Metabolomics & Lipidomics

#### Sample Preparation Plasma

Metabolites and complex lipids were extracted using a biphasic separation with cold methyl tert-butyl ether (MTBE), methanol, and water in deep well plate format. Briefly, 1 ml of ice-cold MTBE and 260 μl methanol was added to 40μl of the plasma spiked-in with 40 μl deuterated lipid internal standards (Sciex, cat# 5040156). The samples were then agitated at 4°C for 30 minutes. After addition of 250 μl of ice-cold water, the samples were vortexed for 1 minute and centrifuged at 3,800 g for 5 minutes at 4°C. The upper organic phase contained the lipids, the lower aqueous phase contained the metabolites, while the proteins were precipitated at the bottom of each well. For quality control purposes, 3 reference plasma samples (40 μl volumes), in addition to one control sample lacking any sample, were processed in parallel per plate. For metabolite preparation, proteins were further precipitated by adding 500 μl of 1:1:1 acetone: acetonitrile: methanol spiked-in with 15 labeled metabolite internal standards to 300 μl of the aqueous phase and 200 μl of the lipid phase, and incubated overnight at -20°C. After centrifugation at 3,800 g for 10 minutes at 4°C, the metabolic extracts were dried down to completion and resuspended in 200 μl 50/50 methanol/water for Liquid Chromatography–Mass Spectrometry (LC-MS) analysis. We examined distribution of metabolites expression and removed one outlier sample from decedent 2 due to poor data quality (**Supplementary Figure 3**).

#### Sample Preparation for Targeted Lipidomics

Lipid extracts were analyzed a Sciex platform that comprises a 5500 QTRAP system equipped with a SelexION differential mobility spectrometry (DMS) interface (Sciex) and a high flow LC-30AD solvent delivery unit (Shimazdu, MD, USA). Briefly, lipid molecular species were identified and quantified using multiple reaction monitoring (MRM) and positive/negative ionization switching. Two acquisition methods were employed covering 13 lipid classes; *Method 1* had SelexION voltages turned on, while *Method 2* had SelexION voltages turned off. High data quality was ensured by i) tuning the DMS compensation voltages using a set of lipid standards (cat# 5040141, Sciex) after either each cleaning procedure, more than 24 hours of standing idle, or 3 days of consecutive use, ii) performing a quick system suitability test (QSST) (catalogue # 5040407, Sciex) before each batch to ensure an acceptable limit of detection for each lipid class, and iii) triplicate injection of lipids extracted from a reference plasma sample (catalogue # 4386703, Sciex) at the beginning of each batch.

#### Data acquisition for Plasma Metabolites

Metabolite extracts were analyzed using a broad-spectrum untargeted LC-MS platform as previously described^21^ while complex lipids were quantified using a targeted MS-based approach^22^.

Untargeted Metabolomics by Liquid Chromatography (LC)-MS: Metabolic extracts were analyzed in quadruplicate using HILIC and RPLC separation in both positive and negative ionization modes. Data were acquired on a Thermo Q Exactive HF mass spectrometer for HILIC (Thermo Fisher, Bremen, Germany) and a Thermo Q Exactive mass spectrometer for RPLC (Thermo Fisher, Bremen, Germany). Both instruments were equipped with a HESI-II probe and operated in full MS scan mode. MS/MS data were acquired on quality control samples (QC) consisting of an equimolar mixture of all samples in the study (global reference pool). HILIC experiments were performed using a ZIC-HILIC column 2.1 × 100 mm, 3.5 μm, 200Å (Merck Millipore, Darmstadt, Germany) and mobile phase solvents consisting of 10 mM ammonium acetate in 50/50 acetonitrile/water (A) and 10 mM ammonium acetate in 95/5 acetonitrile/water (B). RPLC experiments were performed using a Zorbax SBaq column 2.1 × 50 mm, 1.7 μm, 100Å (Agilent, CA) and mobile phase solvents consisting of 0.06% acetic acid in water (A) and 0.06% acetic acid in methanol (B). Data quality was ensured by (i) injecting 6 and 12 pool samples to equilibrate the LC-MS system prior to running the sequence for RPLC and HILIC, respectively, (ii) injecting a pool sample every 10 injections to control for signal deviation with time, and (iii) checking mass accuracy, retention time and peak shape of internal standards in each sample.

#### Data acquisition for Lipidomics

Lipidomics data were reported by Shotgun Lipidomic Assistant (SLA V1.21,^23^ software which calculates concentrations for each detected lipid as average intensity of the analyte MRM/average intensity of the most structurally similar internal standard (IS) MRM multiplied by its concentration. Lipids detected in less than 2/3 of the samples were discarded and missing values were imputed by drawing from a random distribution of low values class-wise in the corresponding sample. Lipid abundances were reported as concentrations in nmol/g.

#### Data Processing for Plasma metabolomics

Data from each mode were independently analyzed using Progenesis QI software (v2.3) (Nonlinear Dynamics, Durham, NC). Metabolic features from blanks and those that did not show sufficient linearity upon dilution in QC samples (r<0.6) were discarded. Only metabolic features present in >2/3 of the samples were kept for downstream analyses. Missing values were imputed by drawing from a random distribution of low values in the corresponding sample. Intensity drift was corrected. Data from each mode were merged and metabolic features were annotated as follows. Peak annotation was first performed by matching experimental m/z, retention time, and MS/MS spectra to an in-house library of analytical-grade standards. Remaining peaks were identified by matching experimental m/z and fragmentation spectra to publicly available databases including HMDB, MoNA, MassBank, METLIN, and NIST using the R package ‘metID’ (v0.2.0). We used the Metabolomics Standards Initiative (MSI) level of confidence to grade metabolite annotation confidence (levels 1-3). Level 1 represents formal identifications where the biological signal matches accurate mass, retention time, and fragmentation spectra of an authentic standard run on the same platform. For level 2 identification, the biological signal matches accurate mass and fragmentation spectra available in one of the public databases listed above. Level 3 represents putative identifications that are the most likely name based on previous knowledge. Metabolite abundances were reported as spectral counts.

### Differential expression analysis for proteomics, metabolomics and lipidomics

Data quality was first examined using distribution plot. We ran PCA to identify potential batch effect. For normalization, we scaled the values to have the same median value using *normalizeMedianValues* function from the *limma* package (version 3.54.2) followed by log2 transformation. We only considered features with less than 50% missing values for the downstream analysis. To assess differential expression, we first grouped time points into early (6 and 12h), mid (18 to 36h) and late stages (after 36h). We then performed differential expression analysis using a linear model (∼ individual + stage) followed by empirical Bayes moderation with the *limma* package.

### Integrative analysis methodology

Processed bulk RNA-seq (PBMC), and cytokine, metabolomic, lipidomic and proteomic data from plasma were normalized and scaled to z-scores. Integrative analysis was performed using *fuzzy c-means clustering* to identify multi-omic analytes that had similar temporal expression patterns across the heart transplant time course. This was performed for each xenotransplant time course independently as well as for combined data. We manually evaluated the downstream multiomic clusters for trends that were correlated with clinical events in the time course, and relevant clusters were selected for further analysis. To test for pathways enriched in the individual clusters, we performed pathway analysis using *Metaboanalyst 5*.*0* (using the *joint protein/metabolite analysis* function).

## RESULTS

### Individual bulk RNA-seq, proteomic, lipidomic and metabolomic datasets

Comparative temporal differential expression analysis of bulk RNA-seq, proteomics, lipidomics and metabolomics for blood samples of two pig heart to human xenotransplants was performed spanning all timepoints described in the Methods section. No significant batch effects were found using PCA analysis for each of the proteomics, lipidomics and metabolomics datasets. **Figure 1** illustrates the comparative temporal differential expression analyses (DEA) individually for each of the bulk RNA-seq (PBMC), proteomic (plasma), lipidomic (plasma) and metabolomic (plasma) datasets. The early phase encompasses the 6hr and 12hr post-transplant timepoints, the mid-phase includes 18 h to 36 h post-transplant, while the late phase comprises all timepoints beyond 36 h.

**Figure 1:**
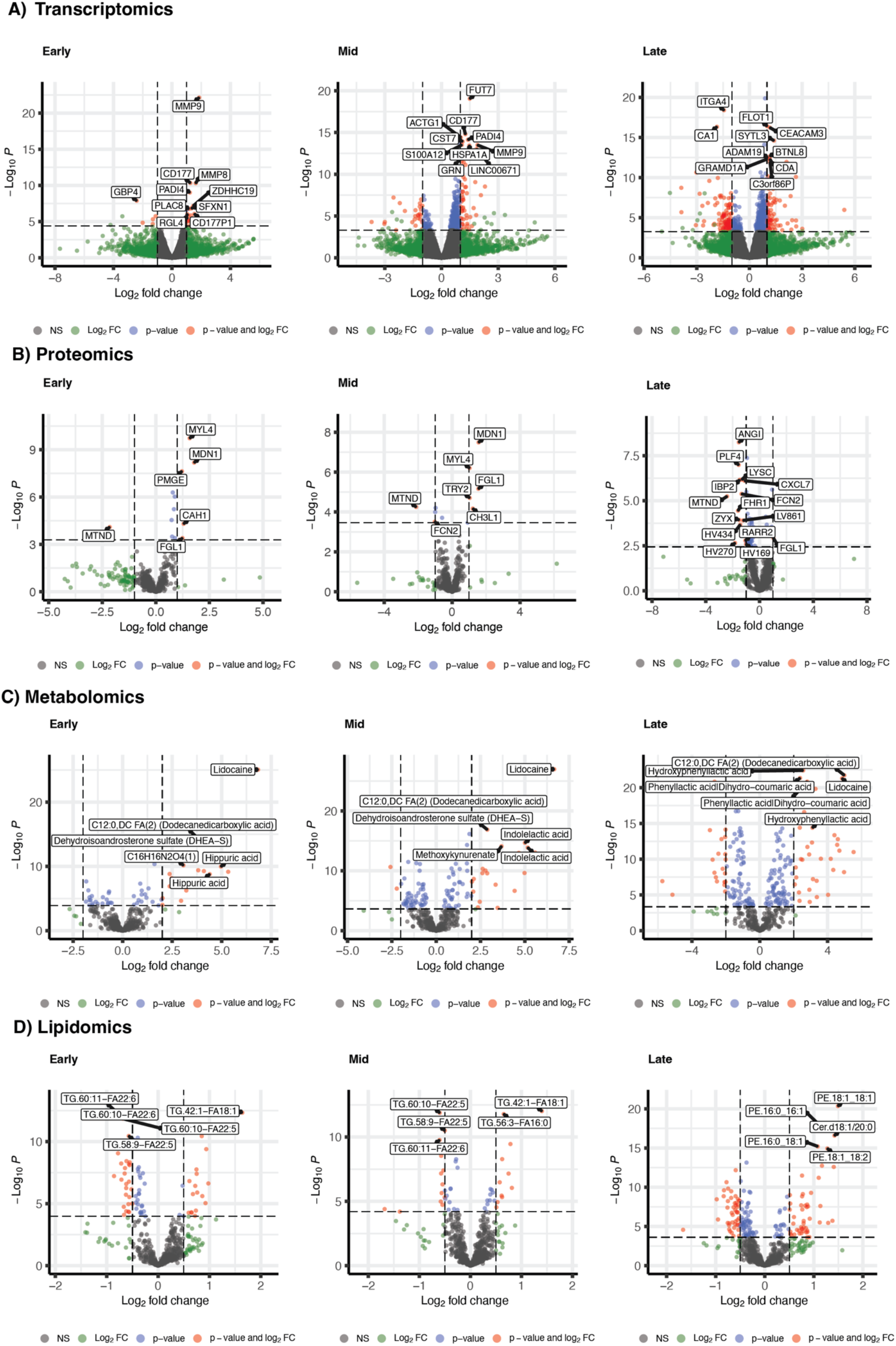
Comparative temporal differential expression analysis of individual omics analyses. Bulk transcriptomics, proteomics, lipidomics and metabolomics for blood samples of two pig heart to human xenotransplantation spanning 26 timepoints are displayed. The early phase encompasses the 6 h and 12 h post-transplant timepoints, the mid-phase includes 18 h to 36 h post-transplant, while the late phase comprises all timepoints beyond 36h. For each panel, the x-axis represents the fold change in log2 scale, and the y-axis depicts the -log10 transformed unadjusted p-value. The individual omics are as follows: **(A) Transcriptomics:** DEA analysis using *DESeq2* is limited to genes whose average expression exceeds 4 counts across samples. Genes with significant expression changes (FDR<0.01) are highlighted in red (absolute log2FC>1) or blue (logFC below cutoff). Only the top 10 most significant genes were labeled due to space limitation. **(B) Proteomics**: The analysis was limited to proteins present in more than half of the samples, yielding a total of 895 proteins. Proteins with significant changes (FDR < 0.05) are highlighted in red (absolute log2FC > 1) or blue (logFC below cutoff). **(C) Metabolomics**: DEA was performed for 459 metabolites. Metabolites with significant changes (FDR < 0.001) are marked in red (absolute log2FC > 2) or blue (logFC below cutoff). **(D) Lipidomics:** The analysis included 720 lipids. Lipids showing significant changes (FDR < 0.001) are denoted in red (absolute log2FC > 0.5) or blue (logFC below cutoff). In the metabolomics and lipidomics panels, only the top 5 features are highlighted with the molecule names. In all panels, molecules without significant changes are represented in gray.

### scRNA-seq of longitudinal PBMCs from two human decedents receiving a pig xenograft heart

We integrated the 26 scRNA-seq samples (14 timepoints for decedent 1 and 12 for decedent 2), to one-dimension reduction space, and used the resulting Leiden clusters to assign PBMC cell-types (**Figure 2A**), with the dynamic proportion of these cell-types shown in **Figure 2B**. We confirmed our cell-type annotations through marker gene discovery (**Figure 2C**) and subsequent gene set enrichment analysis. Interesting trends for cell-type proportions can be observed, with a strong increase of T and NK cells in the later time-points of decedent 1, which is accompanied by erythroblasts, and to a lesser extent, granulocytes. Interestingly, this is at the expense of monocyte proportions. A short-lived increase in the B cell population can be seen in decedent 1 (**Figure 1B**). **Supplementary Figure 4** shows a comparison of the bulk-RNA-seq v<u>ersu</u>s scRNA-seq dataset for specific T and B cells markers. The scRNA-seq cell proportion trends in decedents 1 and 2 are confirmed through analysis of PBMC Bulk RNA-seq (sum of their markers).

**Figure 2:**
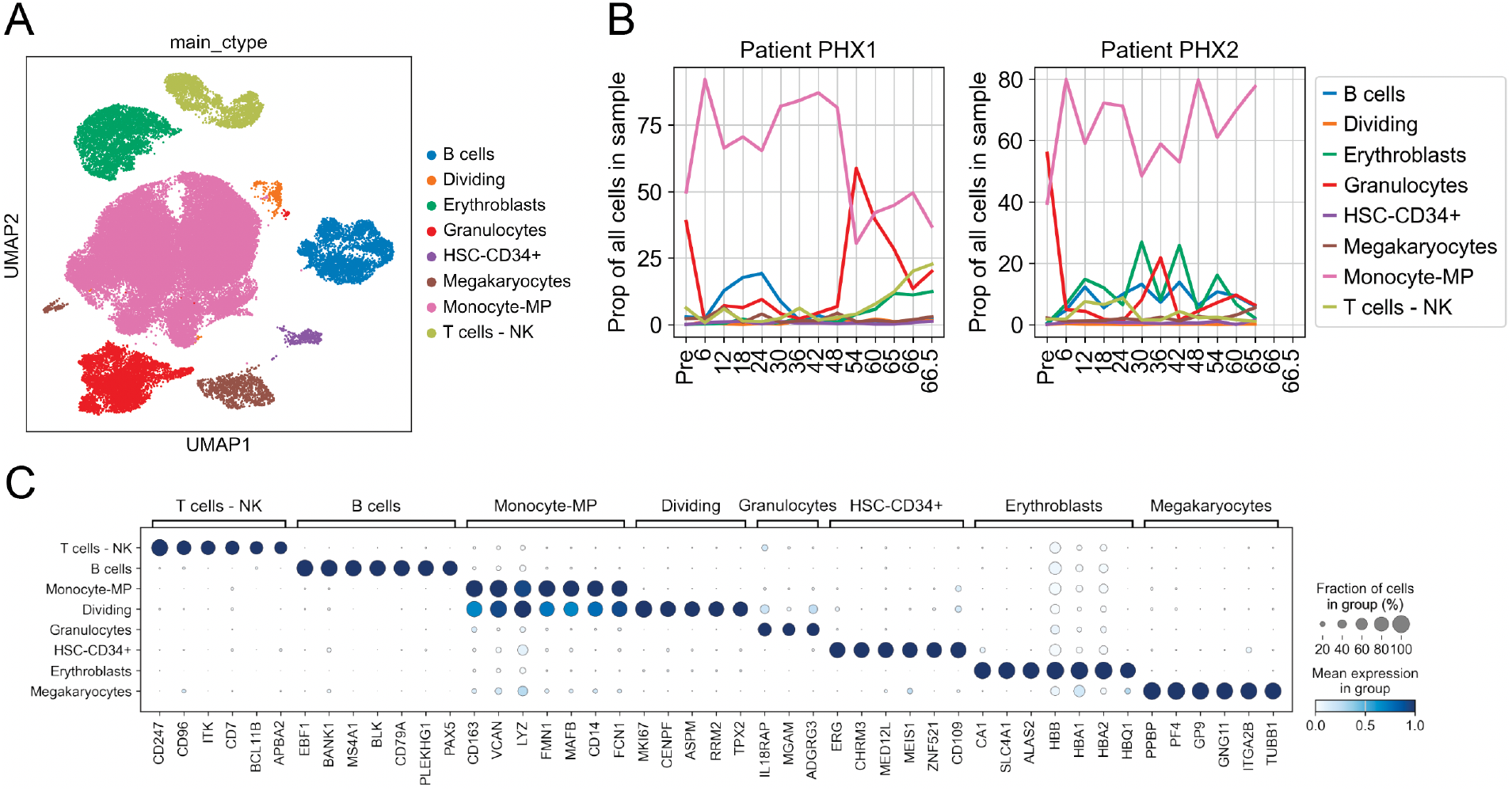
Cell-type diversity from PBMC scRNA-seq, across timepoints and patients. A. Low dimension embedding representation of the PBMC scRNA-seq data, colored by cell-types. B. Proportion (percentages) of each cell-type between samples/timepoints for patient 1 (left) and patient 2 (right). D. Expression distribution of selected marker genes of the main cell-types, basis for their identification.

#### Thymoglobulin immunosuppression treatment dynamics of T-cell depletion in Decedent 1 and 2

Induction immunosuppression for both decedents included rabbit anti-thymocyte globulin (rATG), methylprednisolone, mycophenolate, and eculizumab. Both decedents received methylprednisolone 1,000 mg and rATG 1.5 mg/kg preoperatively; the first decedent received a single dose of rATG 1.5 mg/kg,while the second decedent received an additional dose of rATG on postoperative day (POD) 24.5 hours post-transplant. Both decedents received eculizumab 1,200 mg on POD 0 and 900 mg on POD 1. Maintenance immunosuppression consisted of administering 1,000 mg of methylprednisolone daily, 1,000 mg of intravenous mycophenolate mofetil twice daily beginning POD 0, which continued until the heart was explanted from the decedent at 66 hours post-reperfusion. **Figure 3A&B** illustrates the proportion of T-cell and Natural Killer (NK) cell subtypes overall and across the individual decedent time-courses (the specific marker expression for each cell type is shown in **Figure 3C**). A strong blunting of the all the T-cell subtypes down to negligible readings is evident in decedent 2, after the 24.5 hr post-transplant rATG infusion versus decedent 1 (who did not receive a second rATG dose), and who was observed to have had a pronounced increase in CD4 and CD8 T-cells (**Figure 3b**). We also deconvolved B-cell subtypes across the two xenotransplant time courses (**Figure 4A-C**). Corresponding to the overall increase in B-cells (**Figure 1B**) we observed that this was largely comprised of B-cells expressing TCL1, a marker that is commonly associated with immune tolerance, and to a lesser degree CD70+ B cells. We base cell-type and subtype labelling on marker genes (**Figure 2C, 3C, 4C**) and marker-based gene set enrichment analysis (**Supplemental Figure 5**). A more detailed look into subtype proportion changes (**Supplemental Figure 6**) highlights increase of dividing T cells, FOS+ T cells, Tregs in the later timepoints of patient 1, specifically, in addition of a Plasma cell spike going along the B cell spike in patient 1.

**Figure 3:**
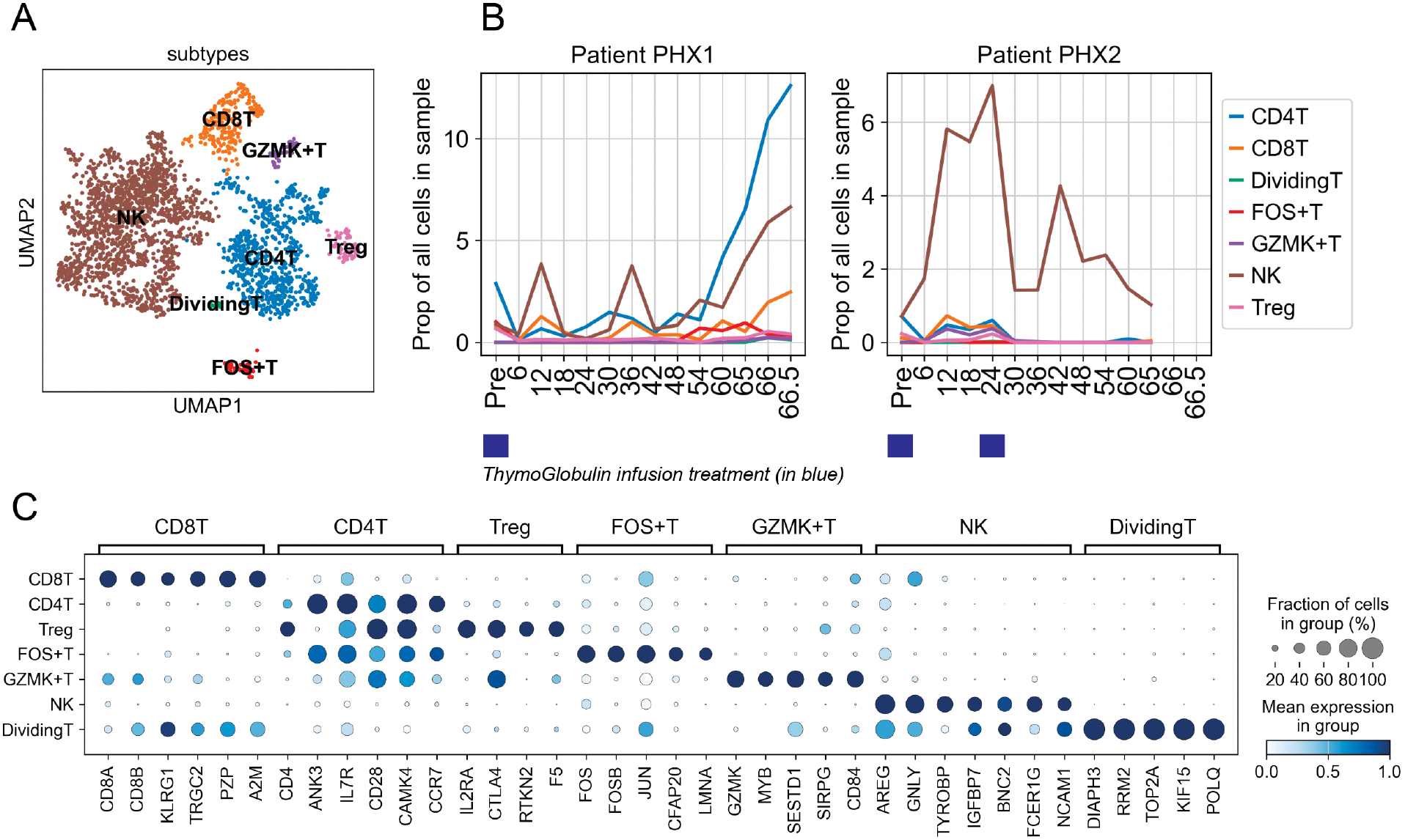
T and NK subtype diversity from PBMC scRNA-seq, across timepoints and patients. A. Low dimension embedding representation of the T/NK cells, colored by subtype. B. Proportion (percentages) of each subtype between samples/timepoints for patient 1 (left) and patient 2 (right). The value represents percentage of the cell-type among all cells of the sample. D. Expression distribution of selected marker genes of the subtype, basis for their identification.

**Figure 4:**
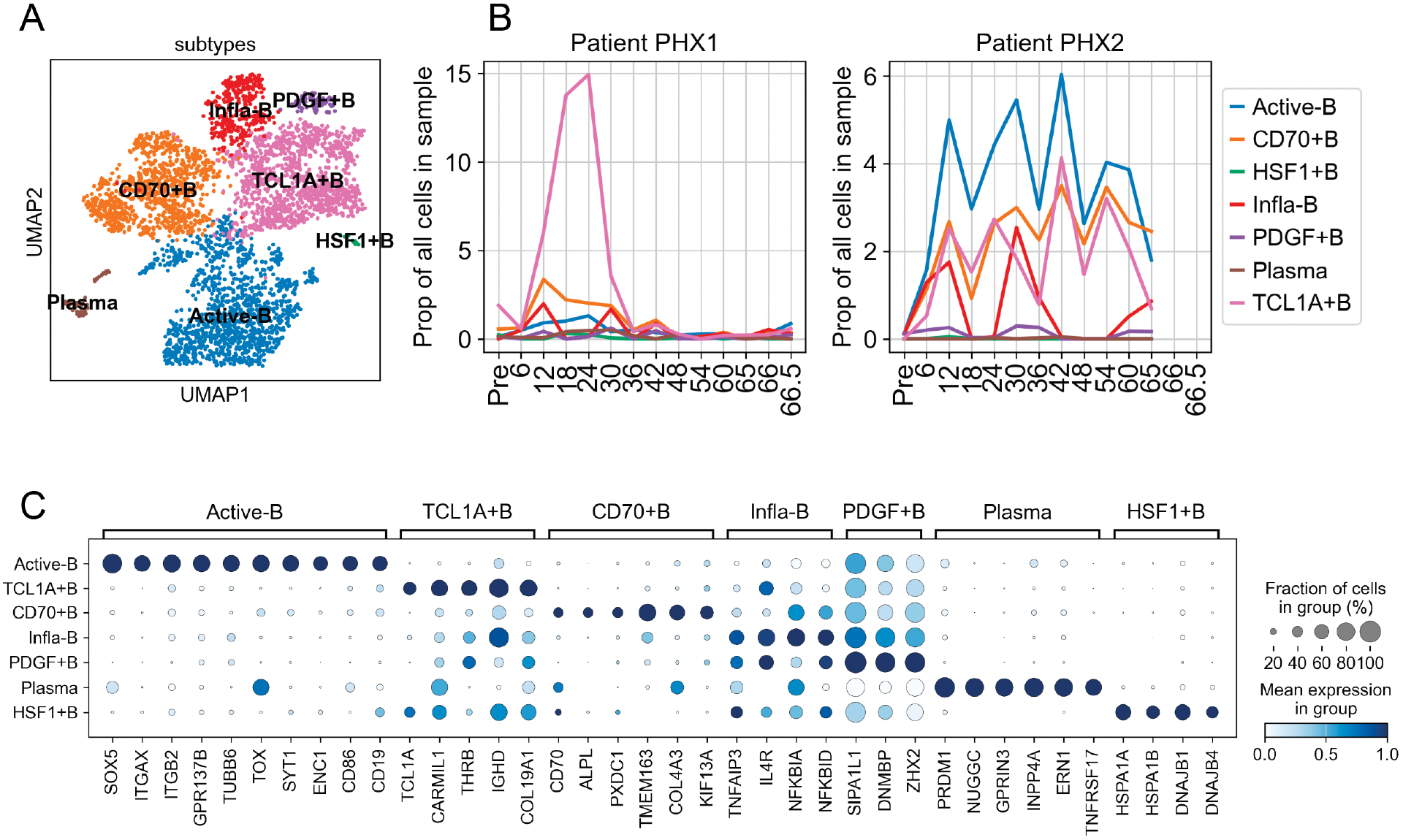
B cell subtype diversity from PBMC scRNA-seq, across timepoints and patients. A. Low dimension embedding representation of the B cells, colored by subtype. B. Proportion (percentages) of each subtype between samples/timepoints for patient 1 (left) and patient 2 (right). The value represents of the cell-type among all cells of the sample. D. Expression distribution of selected marker genes of the subtype, basis for their identification.

### snRNA-seq of pig xenograft heart tissue harvested from human decedent #1 at day 3

In order to compare the immune response in the blood and tissue, we next performed tissue-specific single-nuclei RNA-seq (snRNA-seq) (10x Genomics) using flash frozen and isolated nuclei from 4 × 50uM slices from right ventricle pig heart xenograft explant tissue at the 66 h timepoint. We used the mixed genome alignment method of *CellRanger* to assign a human/pig label to each nucleus and subsequently removed multiplets. We then computationally isolated the nuclei between the pig and human species so they could be analyzed independently. We observed that the human nuclei express macrophages and T cells marker genes (**Supplementary Figure 7**), which is evidence of human immune cells infiltrating into the pig heart xenograft. For the mapped pig cells, **Supplementary Figure 8** shows the main cardiac cell-types observed, including cardiomyocytes, vascular endothelial cells (VECs), fibroblasts, immune cells, smooth muscle cells, pericytes and Schwann cells. We also identify myofibroblasts, and two hypoxic populations (mTORC+ and HIF-1+).

### Geospatial transcriptomics of pig heart xenograft tissue from human decedent #1 at day 3

For geospatial transcriptomics we used the same OCT tissue section as the snRNA-seq samples. For processing, duplicate 10μm OCT tissue sections were analyzed using the Visium v1 platform (10x Genomics) as described in detail in the Methods section. We performed a standard clustering of the Visium data from the fresh-frozen capture areas and observed 10 broad specific groups of cells (**Figure 5A & B)**. Cluster 9 expressed inflammatory/immune related markers such as *CCL21, LYVE1*, while cluster 6 markers are almost exclusively human genes (96 out of 100 top markers, based on Wilcoxon ranked-sum test) (**Supplementary Figures 9 & 10**).

**Figure 5:**
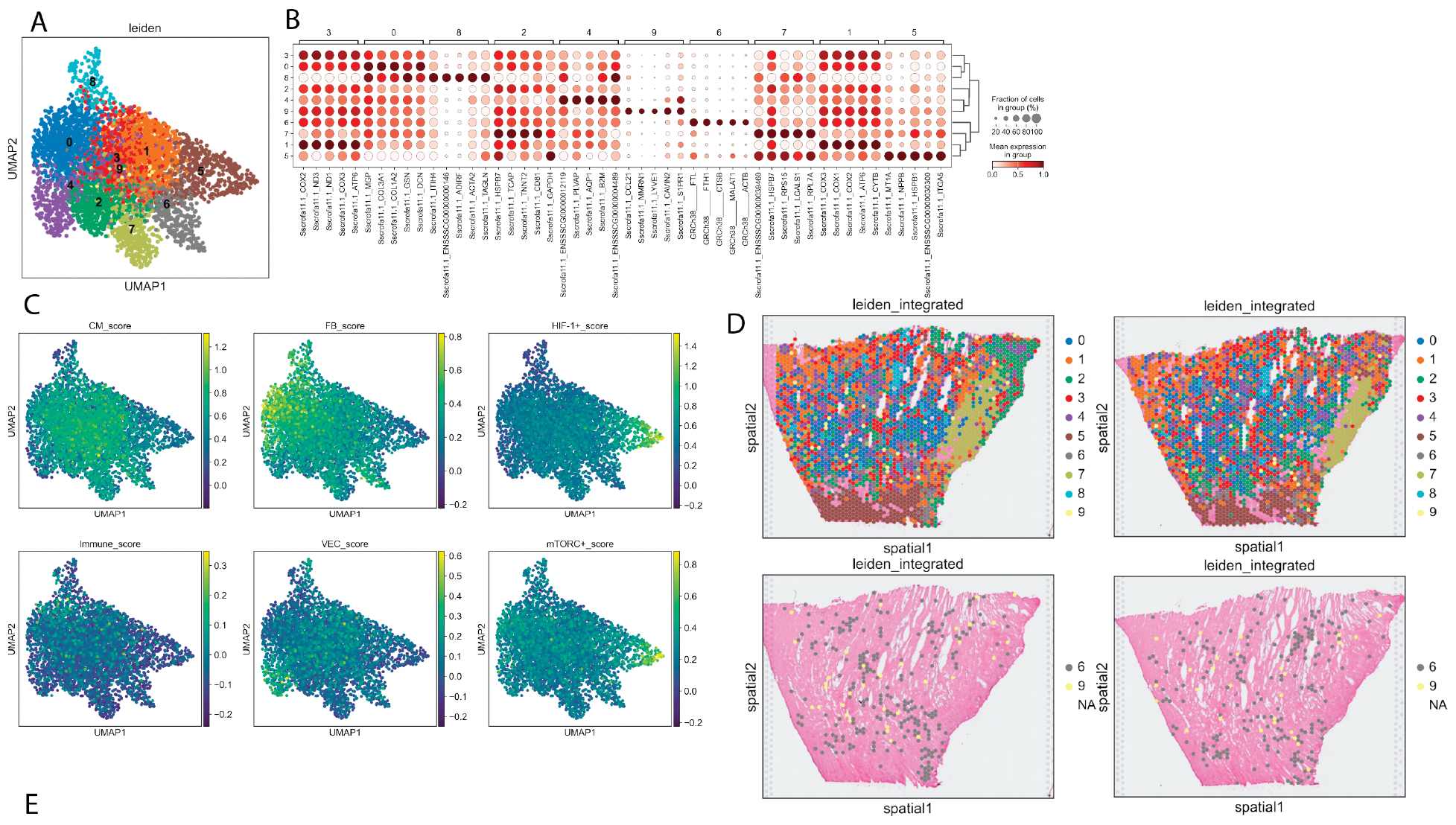
Geospatial transcriptomic analysis of pig and human cells in xenotransplanted tissues. In Visium cluster analysis, ten broad specific capture area groups are identified (A). Cluster 9 expressed inflammatory/immune related markers such as *CCL21, LYVE1*, while cluster 6 markers are almost exclusively human genes (B). Visium data cell-type signatures expression evaluation vs snRNA-seq data (C). We observe that the capture areas in cluster 0 are enriched for fibroblasts, cluster 5 in hypoxic cells, cluster 4 in vascular endothelial cells (VECs). Interestingly, we also see an enrichment of EC and VECs markers in cluster 9, which appears to be associated with inflammatory signals. By mapping the capture areas on the slides at their location, we can assess the spatial distribution of the observed clusters (D).

We then used *scanpy* gene score function to further evaluate cell-type signatures expression on the Visium data, from the signatures observed in the snRNA-seq data (**Figure 5C**). We observe that the capture areas in cluster 0 are enriched for fibroblasts, cluster 5 in hypoxic cells, cluster 4 in vascular endothelial cells (VECs). Interestingly, we also see an enrichment of EC and VECs markers in cluster 9, which are associated with inflammatory signals. By mapping the capture areas on the slides at their location, we can assess the spatial distribution of the observed clusters (**Figure 5D**). Cluster 5 and 7 have a consistent location on the bottom and the right of both slides. When focusing on the human cluster 6 and pig inflammatory 9 clusters, we identify two trends: the human cells are mostly distributed in hotspots and accompanied by inflammatory pig cells.

H&E stains were also assessed by an expert clinical transplant pathologist. **Supplementary Figure 11** (A-C), in decedent 1, shows coagulative necrosis, which may be indicative of interstitial edema and endothelial swelling in a small interstitial capillary. In the same individual endothelial swelling in the intramyocardial muscular arteries and perivascular edema is evident along with a ‘lifting’ effect with of detached endothelial layers, although few inflammatory cells are evident. Decedent 2 also has endothelial swelling evident in intramyocardial muscular arteries and small interstitial capillaries Supplementary Figure 11 (D-E).

#### Pathway Analyses

When focusing on the top 100 marker genes of the human cell cluster 6, we aimed to identify these cells and their function in the context of the xenotransplant. We removed the four pig marker genes as well as mitochondrial and ribosomal human genes, and subsequently performed over-representation analyses using *enrichr* and *enrichr-KG* ^24^. The pathway results, summarized in **Supplementary Figure 12** (top 5 gene-sets by p-value of the 4 gene sets libraries), strongly indicate expression patterns consistent with antigen mediated immune response to the graft, with terms such as *Allograft Rejection, Antigen Processing, Neutrophil Degranulation, Macrophage markers*. Other enriched pathway terms such as *VEGFA-VEGFR2 Signaling* and *Viral myocarditis* may present potential interactions between the endothelial cells and cardiomyocytes. The MSigDB Hallmarks gene-sets enrichment analyses confirm these observations with *Allograft Rejection* and *Angiogenesis* being ranked as the 3rd and 4th most significant pathways. To cross-validate the human-cell expression signals from both the snRNA-seq and Visium datasets, we co-visualized the Visium cluster six markers with the human nuclei RNA signal to and confirm their expression in both independent experiments performed on the same tissue (**Supplementary Figure 9**).

#### Endothelial immune response

To echo on the endothelial stress observed on H&E staining (**Supplementary Figure 11**), we assessed the transcriptional identity of endothelial cells from xenoheart snRNA-seq (pig cell selections). We identify two distinct clusters of endothelial cells (EC and VECs, **supplemental figure 8)**. These clusters both display endothelial cell related pathways from their marker genes overrepresentation analysis (GSE, **supplemental figure 13**), in addition to unexpected immune pathways, which are mostly only enriched in the endothelial and immune clusters. These pathways include T cell and B cell receptor pathways. We then seek to validate this immune / EC signal in the Visium spatial transcriptomics data. First, we use gene-set enrichment on Visium cluster markers and show the enrichment of immune related gene sets in cluster 4 (cytokine mediated signaling, neutrophil degranulation, **supplemental figure 14A**). Secondly, we calculate the correlation of lead gene expression score (from the immune EC/pathways of **supplemental figure 13**), and the Visium cluster score. We see positive and significant correlation for most of these pathways in cluster 4 (**supplemental figure 15A-B**), confirming their specificity to that cluster. These analyses enable us to confirm the endothelial label of Visium cluster 4, through enrichment of cluster 4 gene score with EC / VEC marker gene scores (**supplemental figure 15C-D**), and through the GSE analysis of Visium cluster 4 marker genes (positive regulation of angiogenesis, positive regulation of vasculature development, **supplemental figure 14A**). Finally, we see a positive link between the EC / immune signal found in the snRNA-seq, and confirmed in the Visium data, with the presence of human cell (**supplemental figure 16**). Correlation between the pathway score of each capture area and their fraction of counts aligning to the human genome, partitioned by Visium clusters, is always positive when significant. Moreover, Visium cluster 4 shows the higher number of significant positive correlation out of all Visium clusters.

### Pig to human Integration of bulk RNA-seq, proteomic, lipidomic and metabolomic PBMC datasets

For the PBMC omics datasets, bulk RNA-seq, proteomic, lipidomic and metabolomic datasets from all of the same 6-hour timepoints were integrated into a longitudinal analysis to identify multi-omic signatures associated with post-xenotransplant phenotypes and outcomes. Omics data from timepoints spanning every 6 hours from pre-transplant (0hrs) until 66 hours in xenoheart decedent 1 and 2 were processed with no significant batch effects were found using PCA analysis for each of the single omic procedures. The resultant datasets were integrated into a longitudinal analysis to identify multiomic signatures associated with time post-xenotransplant and other clinical features, using fuzzy c-means clustering after normalizing and standardizing the bulk RNA-seq, proteomic, lipidomic and metabolomic datasets into a unified integrative data matrix^25^. Fuzzy c-means clustering showed a number of clear patterns of analyte abundance changes over the post-xenotransplant monitoring period. An example of this is illustrated in **Figure 6** below where, we show common analytes (RNA-seq, proteins, lipids, cytokines and metabolites) that show progressively increasing abundance starting at the 42-hour mark post-transplant specifically in decedent 1, which track consistently with 3 biomarkers shown across all timepoint: the liver function enzymes, alanine transaminase (ALT) and Aspartate transferase (AST) and a metric of clotting time INR. This cluster was highly enriched for metabolic processes (glycolysis q < 5x10^−13^, pyruvate metabolism q < 5x10^−6^) lipid metabolism (peroxisome, fatty acid degradation q < 2x10^−6^) as well as immune response (antigen presentation and processing q < 0.01).

**Figure 6:**
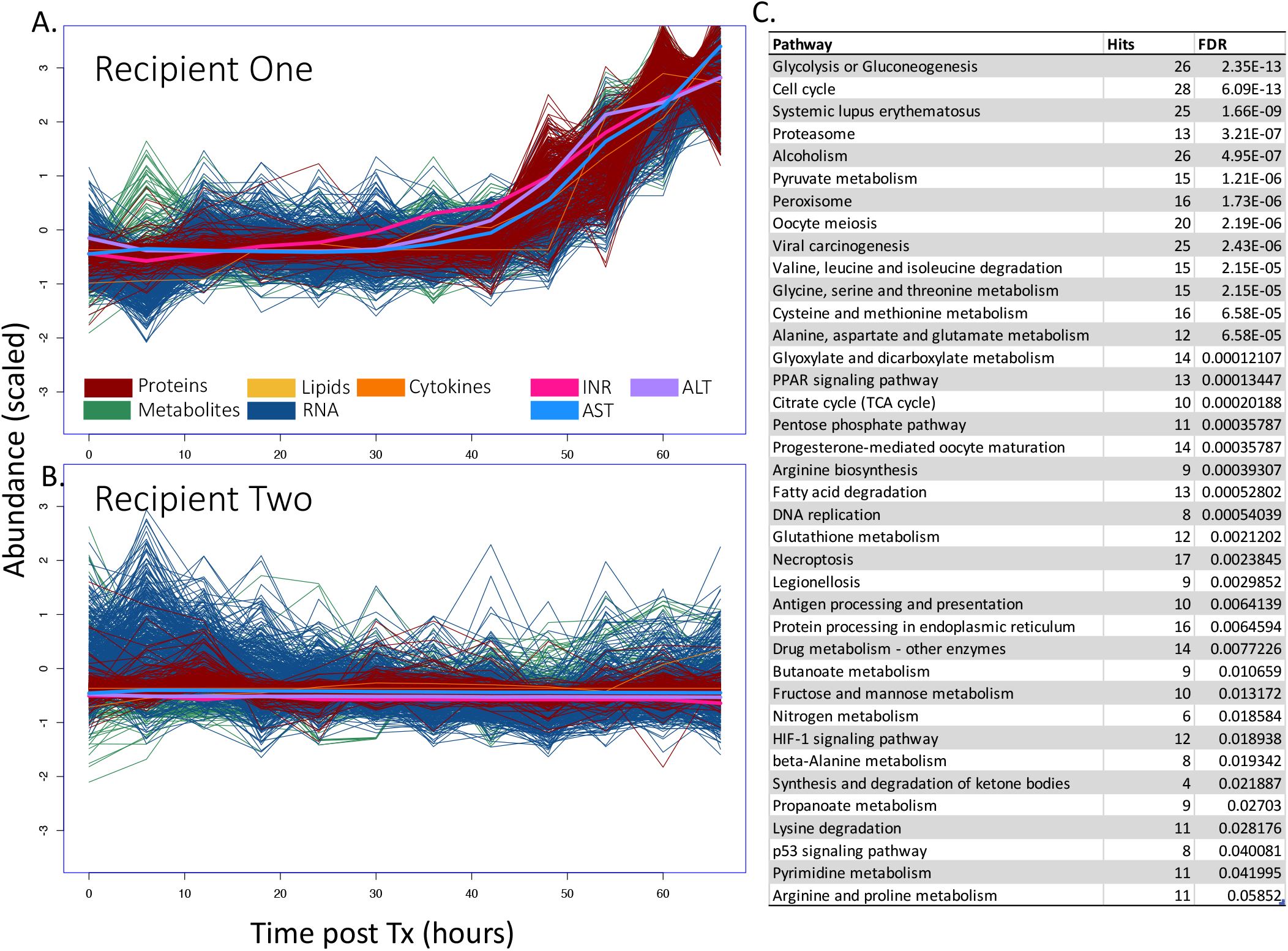
Integrative clustering of transcriptomics, proteomics, metabolomics, lipidomics cytokines from sera of 2 human decedents receiving a pig heart xenograft. Integrative clustering of bulk RNA-seq, proteomics, metabolomics, lipidomics and cytokines using fuzzy c-means clustering across all timepoints reveals an upregulated cluster of analytes at late timepoints that correlated with the indicated clinical variables (INR, AST, ALT). A) shows data for the first xeno timecourse, while B) shows data for the second. The different omics subtypes are color-coded as indicated in the legend. C) shows significant pathways associated with the indicated cluster.

## DISCUSSION

*Sus scrofa domesticus* holds significant promise as a robust source of donor organs that could solve the current lack of availability of human heart allografts. Given the novel setting and lack of *in vivo* data for pig-to-human heart xenotransplantation, comprehensive molecular profiling over time of individuals receiving a pig-derived cardiac transplant will help us better understand how human physiological systems respond to xenotransplantation, and how this response is similar to or different from human-to-human heart allo-transplantation. Here, we have performed the first integrative personal omics profiling (iPOP) of pig-to-human heart xenotransplantation in a human decedent model. Using a dense time course of blood and tissue sampling over a three-day period coupled with large-scale single-cell and bulk multi-omics, we have compiled a detailed portrait of the function and dysfunction of biological systems post xenotransplant.

Overall iPOP profiling shows a markedly different trajectory between the two independent xenotransplant experiments, with decedent 1 showing evidence of dysfunction across many different biomolecule types and scales from about 42 hours onwards, whereas decedent 2 exhibits only minor perturbations throughout the whole procedure. These findings mirrored the clinical course of the two hearts (Moazami et al in press). There are at least 3 possible explanations for the declining function of the first heart: (1) The pig xenograft heart was undersized in the first decedent, resulting in an insufficient volume of blood out being pumped out of the left ventricle during each systolic cardiac contraction; (2) the heart was deteriorating due to early perioperative cardiac xenograft dysfunction (PCXD); (3) post-transplant xenograft dysfunction potentially due to early AbMR from minimized induction immunosuppression protocol or (4) injury from the operative procedure that necessitated placement of bovine pericardiac patches to both the aorta and anterior pulmonary artery in order to bridge size discrepancies between the xenograft heart and the recipient’s vessels.

Due to the novelty of pig-to-human cardiac xenotransplantation, there are no standard guidelines for selecting the appropriately sized xenograft organ for a given recipient’s weight and height. Therefore, the relatively smaller xenograft heart for decedent #1’s size may have catalyzed systemic hypoperfusion and subsequent ischemic injury. In addition, this size mismatch necessitated additional technical steps in the operative procedure for implantation, which may have initiated complement activation and worsened hypoperfusion of the small xenograft heart. The second possibility, PCXD, when a cardiac xenograft shows dysfunction in the first 24-48 hour post-transplant in the absence of rejection^26^, is postulated to be due to a cascade of inflammatory reactions in response to prolonged ischemic time. PCXD is a nonspecific term that includes xenograft deterioration from both poor preservation protection resulting in ischemia reperfusion and immune mediated injury, most often from non-alpha Gal antibodies^26^. Many xenotransplantation researchers have tried to overcome PCXD by modifying myocardial preservation and procurement techniques ^27,28^.

Among the responses occurring in decedent 1, we observed a rapid remodeling of both the tissue immune microenvironment (spatial RNA-seq and snRNA-seq) and in the circulating immune response (scRNA-seq), with a rapid induction of B-cells that tapered off and was later replaced with a robust T-cell-mediated response. iPOP data also showed a sudden increase of CD4+ T cells along with sCD40L about 42hrs post-transplant. This event may have been the beginning of an AbMR process, as CD4+ T cells activate B cells via CD40-CD40L interaction. While a number of the molecular pathway signatures from the PBMC scRNAseq, xenograft snRNAseq and spatial transcriptomics are consistent with molecular AbMR, the conventional histopathology does not show manifestation of these early molecular events. Furthermore, the reactive C4d immunohistochemistry stains in cardiomyocytes confirm myocyte injury may be due to hypoxia and vascular injury, leading to activation of epithelial mesenchymal transition (EMT) and angiogenesis pathways.

The second dose rATG treatment in decedent 2 may be a potential key intervention that explains some of the differences between the dynamic cell-type composition of decedent 1 and 2 over the course of the 3-day study. While the early B-cell memory response at 6-12 hours and subsequent T-cell activation in decedent 1 is observed at ‘n of 1’ experimental level, it is evident that the trajectory is impacted by a second dose of rATG.

In decedent 1, we observed a continually increasing abundance of a large number of omic biomarkers in the blood, starting around the 42 h mark post-transplant. These clusters are highly enriched for metabolic processes (glycolysis, pyruvate metabolism), lipid metabolism (peroxisome, fatty acid degradation), liver injury as well as immune response (antigen presentation and processing) and necroptosis. This mirrors the trajectory of multiple clinical biomarkers for liver dysfunction (prothrombin (INR), alanine transaminase (ALT) and aspartase transaminase (AST)) and likely represents the onset of organ failure and consistent with the detection of necroptosis pathway activation. Given the significant upregulation of glycolysis-related pathways, this may be due to a transition to an acute hypoxic state associated with impaired oxygen profusion from the distressed xenograft and vasopressor use. Of equal importance is the lack of molecular signatures for immune activation or tissue injury in the decedent 2.

While these data represent a novel window into the molecular dynamics during and post xenotransplantation, more study is needed. A key limitation of the current study is that only two xenotransplants were performed and characterized and only over a 3-day period. These 3-day timeframes were initially allowed as this is the typical timeframe that a recently deceased donor with acceptable organs for transplantation is maintained while allocation and procurement proceed. Extending the timeframe of these pig-to-human decedent xenotransplants to a month would add significant value in determining a prolonged period of immune-response and would allow different immunosuppression regimes to be assessed.

It is well established in the xenotransplantation field that costimulatory blockade drugs are essential for the successful protection of xenografts. The absence of costimulatory blockade drugs in the minimized immunosuppression regime of these two human decedents receiving a pig heart xenograft would be considered suboptimal to support long term survival of a xenograft. The longitudinal transcriptomic datasets from the first decedent are likely explained by the lack of costimulatory blockade. It is note that the second treatment of rATG at 24.5 hours in second decedent had a pronounced positive effect on the subsequent immune-cell proportions. In conclusion, we present a wealth of multi-omic data that lays the groundwork for future studies in larger cohorts of humans receiving xenotransplants. Dense longitudinal sampling was necessary for clearly delineating the trajectories of many of these analytes and indicate the value of detailed molecular monitoring to be performed on all future xenotransplantation cases.

