## Supplementary Figures for "Integrative Multi-omic Profiling of Two Human Decedents Receiving Pig Heart Xenografts Reveals Strong Perturbations in Early Immune-Cell and Cellular Metabolism Responses"

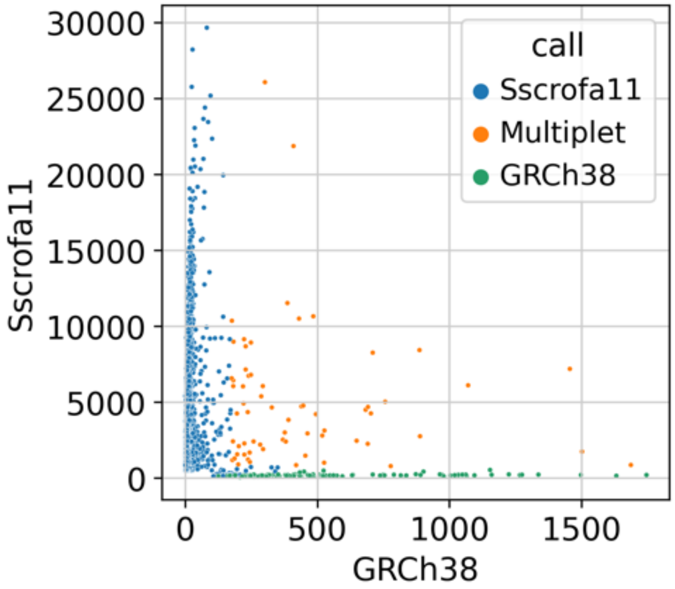


**Supplementary Figure 1:** Distribution of the number of unique molecular indices (UMI) counts mapping to the human (x axis) or pig (y axis) genome from the Xenoheart snRNA-seq, resulting from the *Cellranger* '*Barnyard*' experiment multiple genome aligner. Nuclei are then assigned a genome based on that distribution, and cross-species multiplets are called.

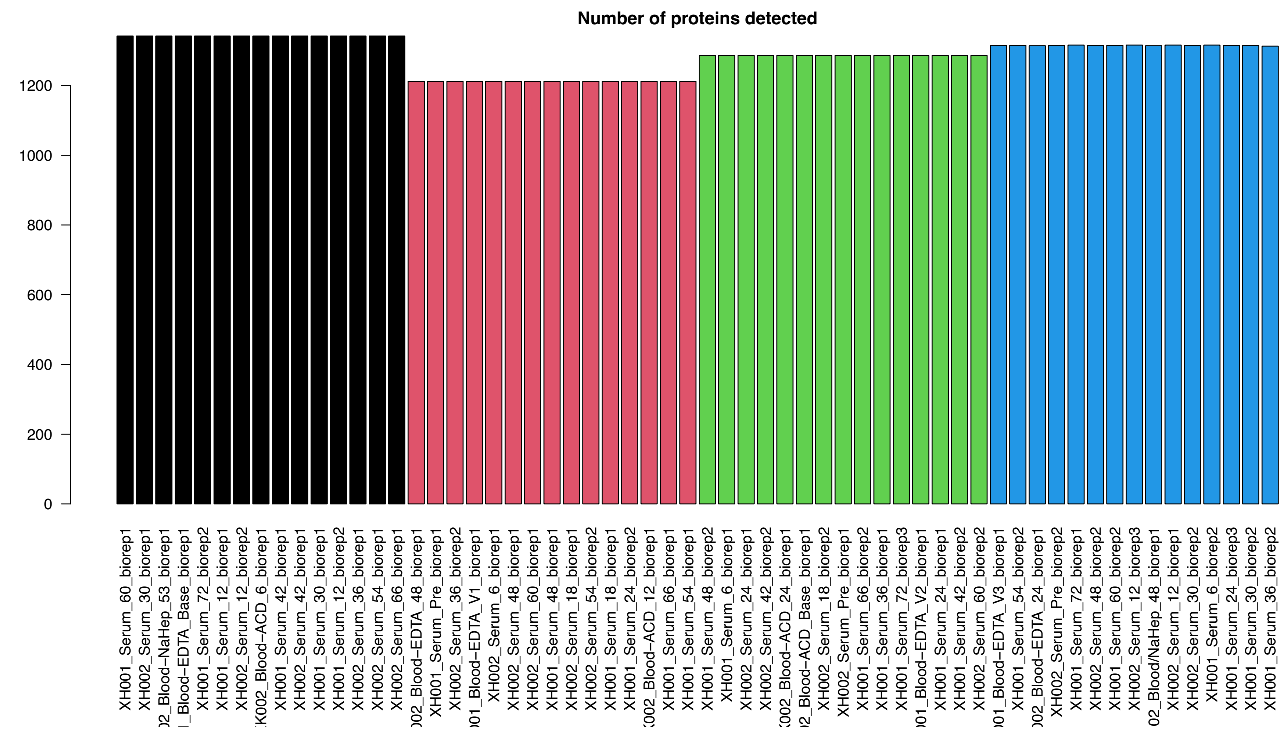


**Supplementary Figure 2: average number of identified protein groups for each sample.** Samples for proteomics were split into 4 run. Over 1000 features were identified for all samples.

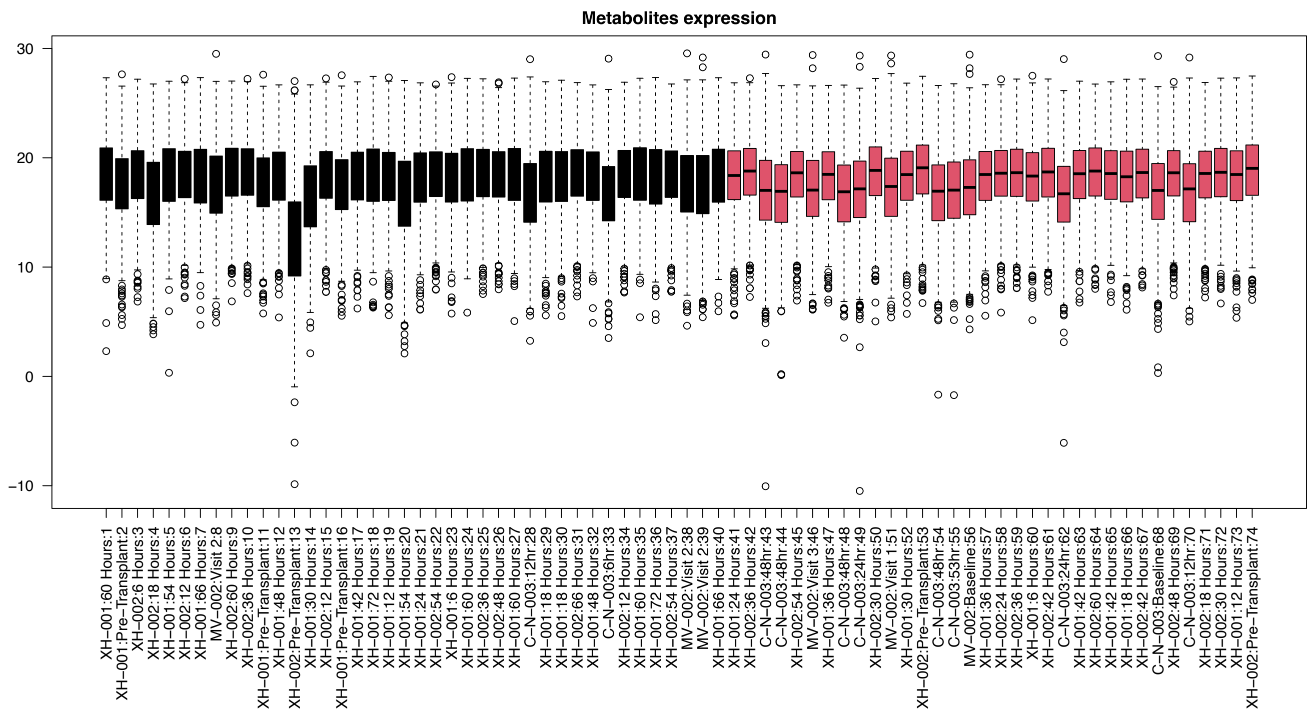


**Supplementary Figure 3: signal distribution for metabolites expression.** One outlier XH-002 (decedent 2):Pre-Transplant:13 was removed for downstream analysis due to poor data quality.

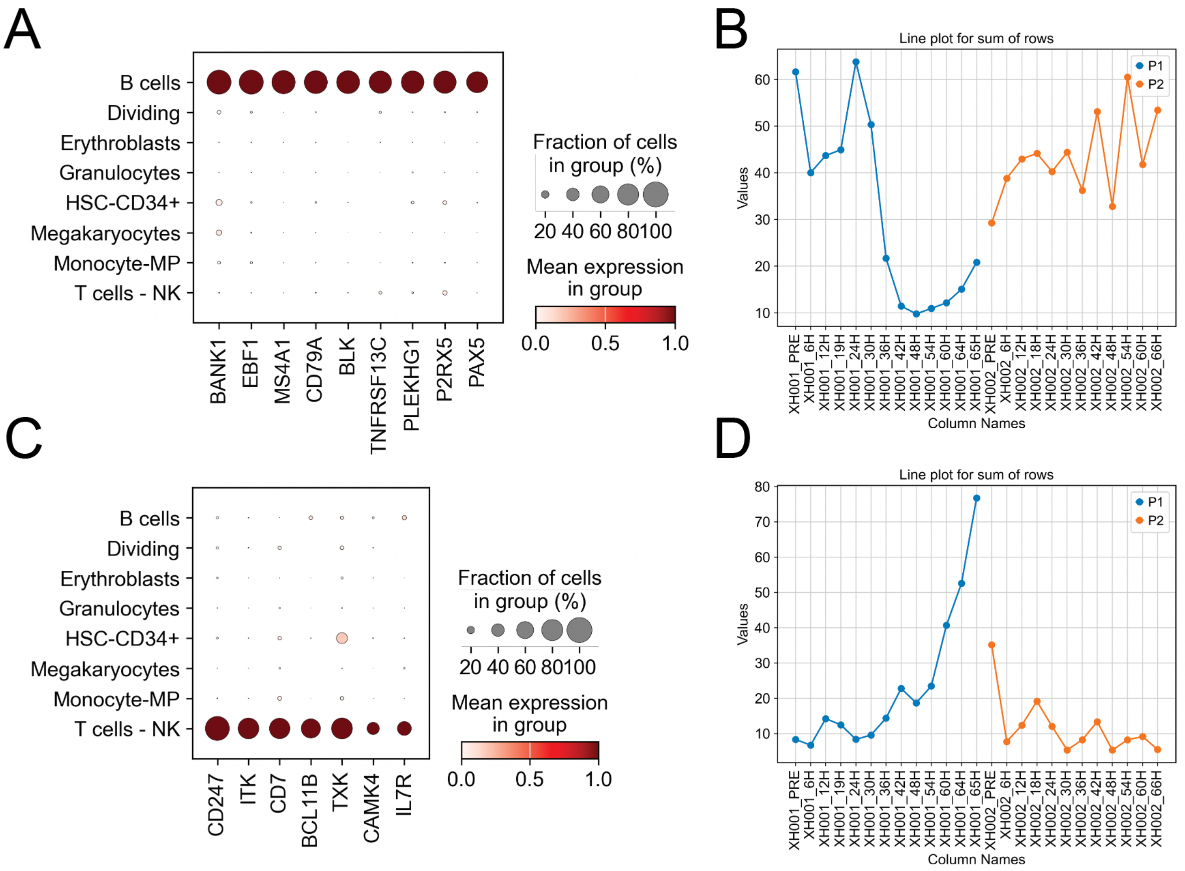


**Supplementary figure 4:** Marker gene expression distribution amongst main cell-types of the PBMC scRNA-seq, for hand selected markers that are specific of B cells (A) and T/NK cells (C). Time-point dependent distribution of the sum of normalized expression of the B cell (B) and T/NK (D) markers from the Bulk-RNA seq data, confirming the proportion trends observed from the scRNA-seq.

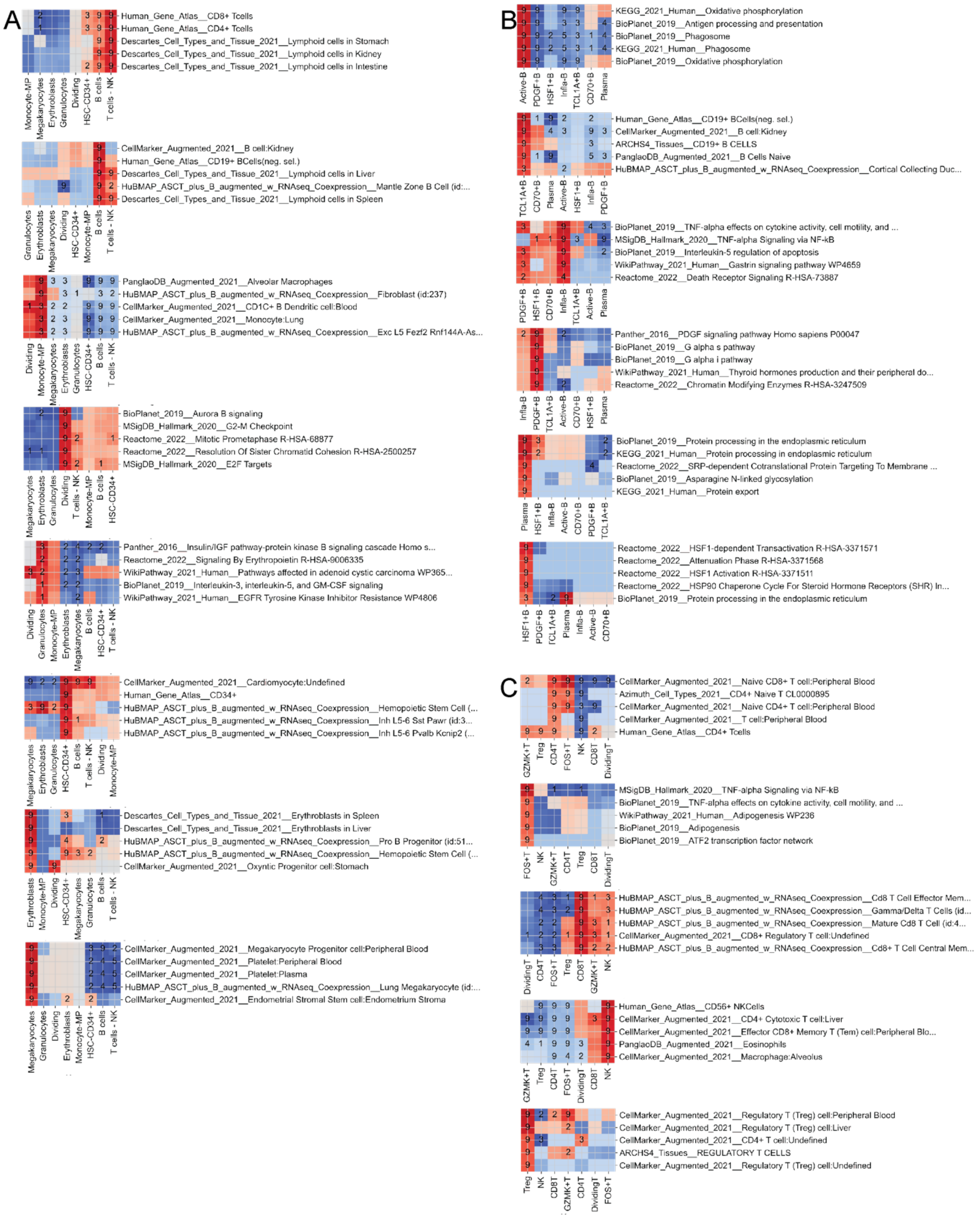


**Supplemental figure 5:** Cell type and subtype markers genes enrichments (GSEA). A. Main cell types. B. B cells. C. NK and T cells.

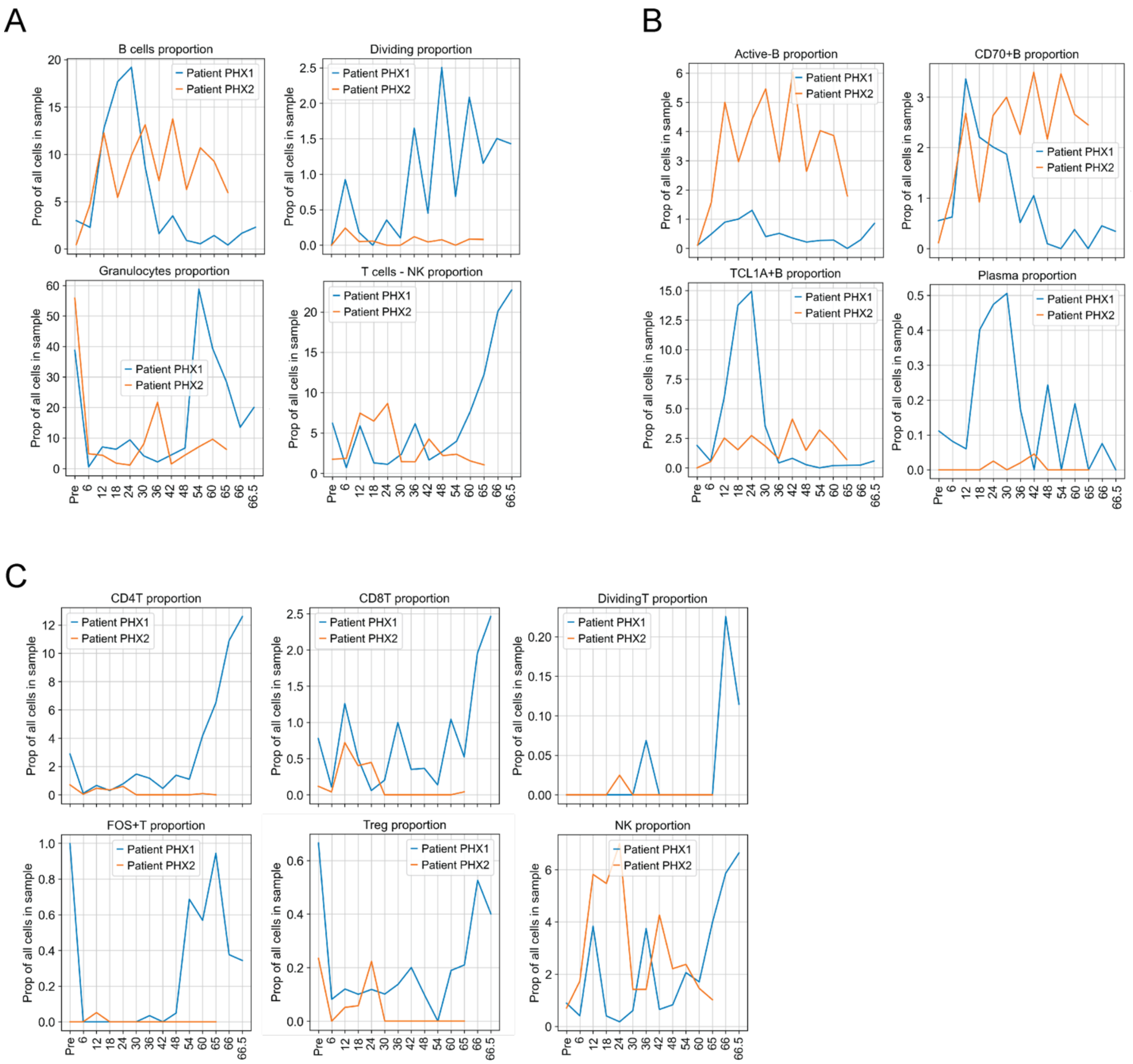


**Supplemental figure 6**: Cell-type and subtype proportion distribution across timepoint and patients. The percentage of presence of that cell-type / subtype among all cells of each sample is represented. This figure shows specific groups that display percentage patterns of interest, focusing on main cell-types (A) B cells subpopulations (B) and T/NK subtypes (C).

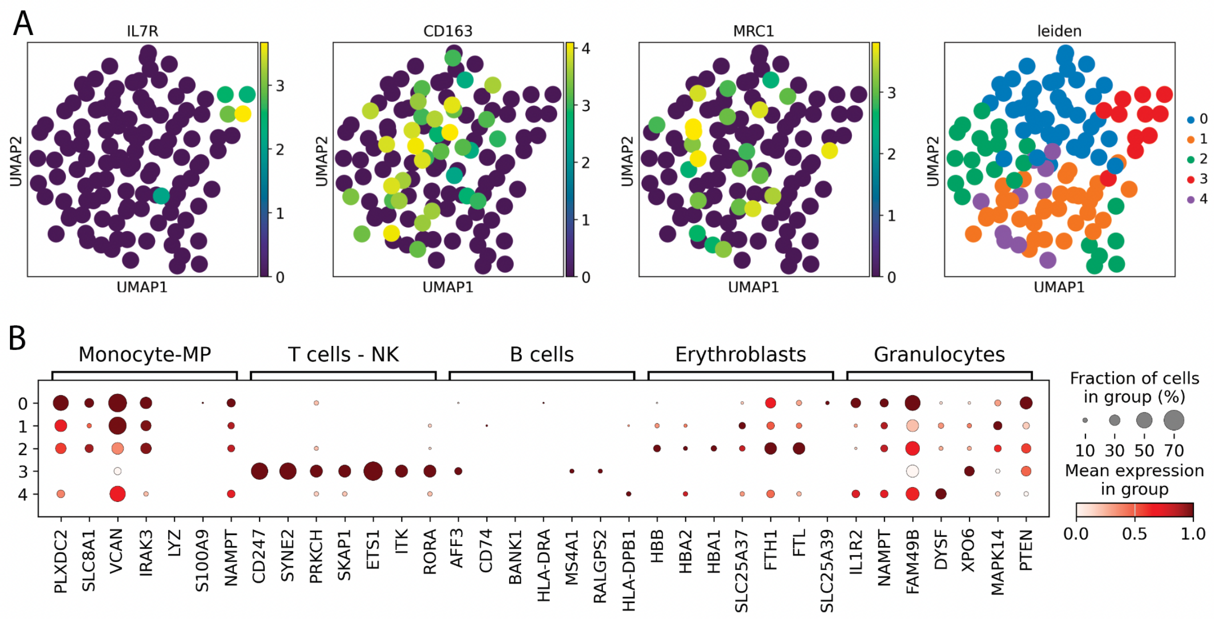


**Supplementary Figure 7:** Heart tissue snRNA from Xenoheart 1 Day 3 timepoints. A. Distribution of the human cell transcriptomic diversity, with key immune marker genes and clusters. B. Expression distribution of top 7 markers of the main PBMC cell-type markers, within the human nuclei present in the Xenoheart.

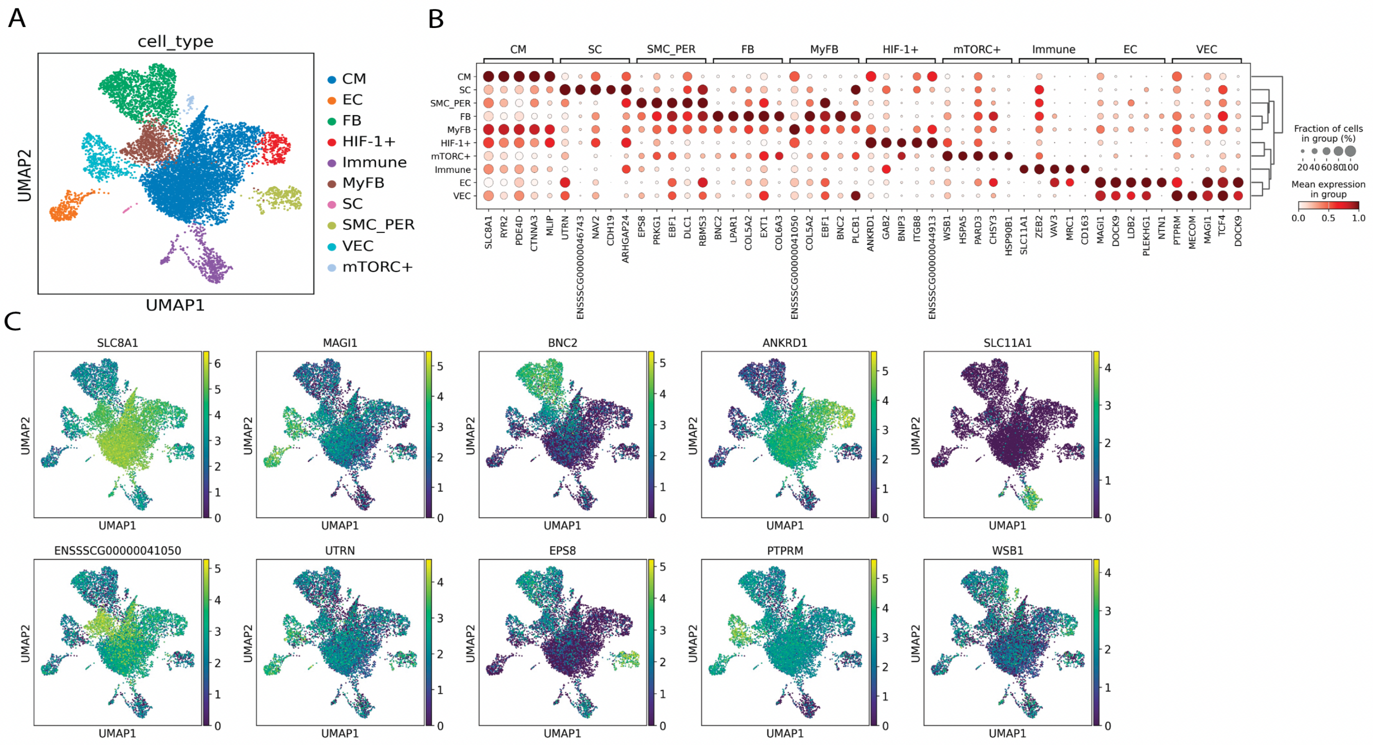


**Supplementary Figure 8:** Main cell-type distribution from Xenoheart pig nuclei (snRNA-seq). A. low dimension embedding distribution of the pig nuclei (Xenoheart snRNA-seq), colored by assigned cell-type. B. Expression distribution of the top 5 differentially expressed genes for each cell-type group (Wilcoxon rank-sum test), showing the main cardiac cell-types observed, including cardiomyocytes, vascular endothelial cells (VECs), fibroblasts, immune cells, smooth muscle cells, pericytes and Schwann cells. Myofibroblasts, and two hypoxic populations (mTORC+ and HIF-1+) are also observed. C. Expression distribution on the UMAP embedding of the top marker for each identified cell-type.

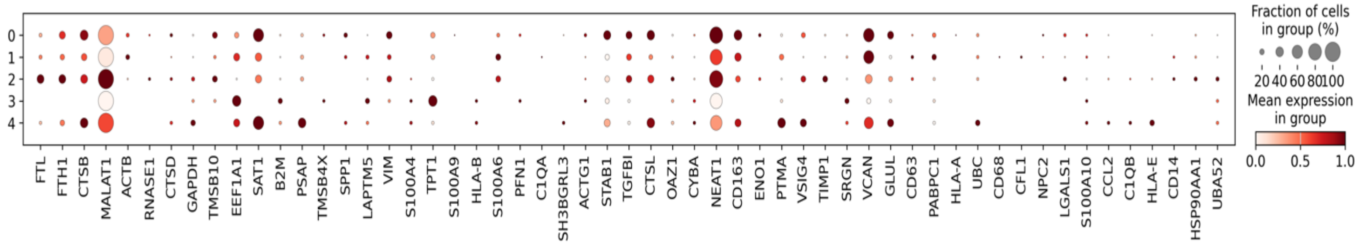


**Supplementary Figure 9**: Expression distribution of the top marker genes of the cluster 6 (human cluster) of the Visium data, among the human nuclei from the pig heart xenograft snRNA-seq clusters. The markers represent the main human genes observed from the Visium assay, confirming their presence.

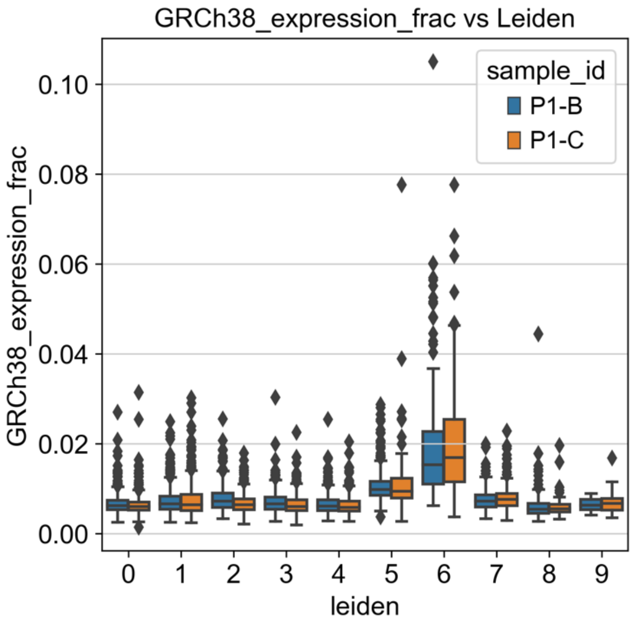


**Supplementary Figure 10:** Cluster 6 is the most enriched cluster for counts aligning to the human genome. Distribution of proportion fraction among the Visium clusters.

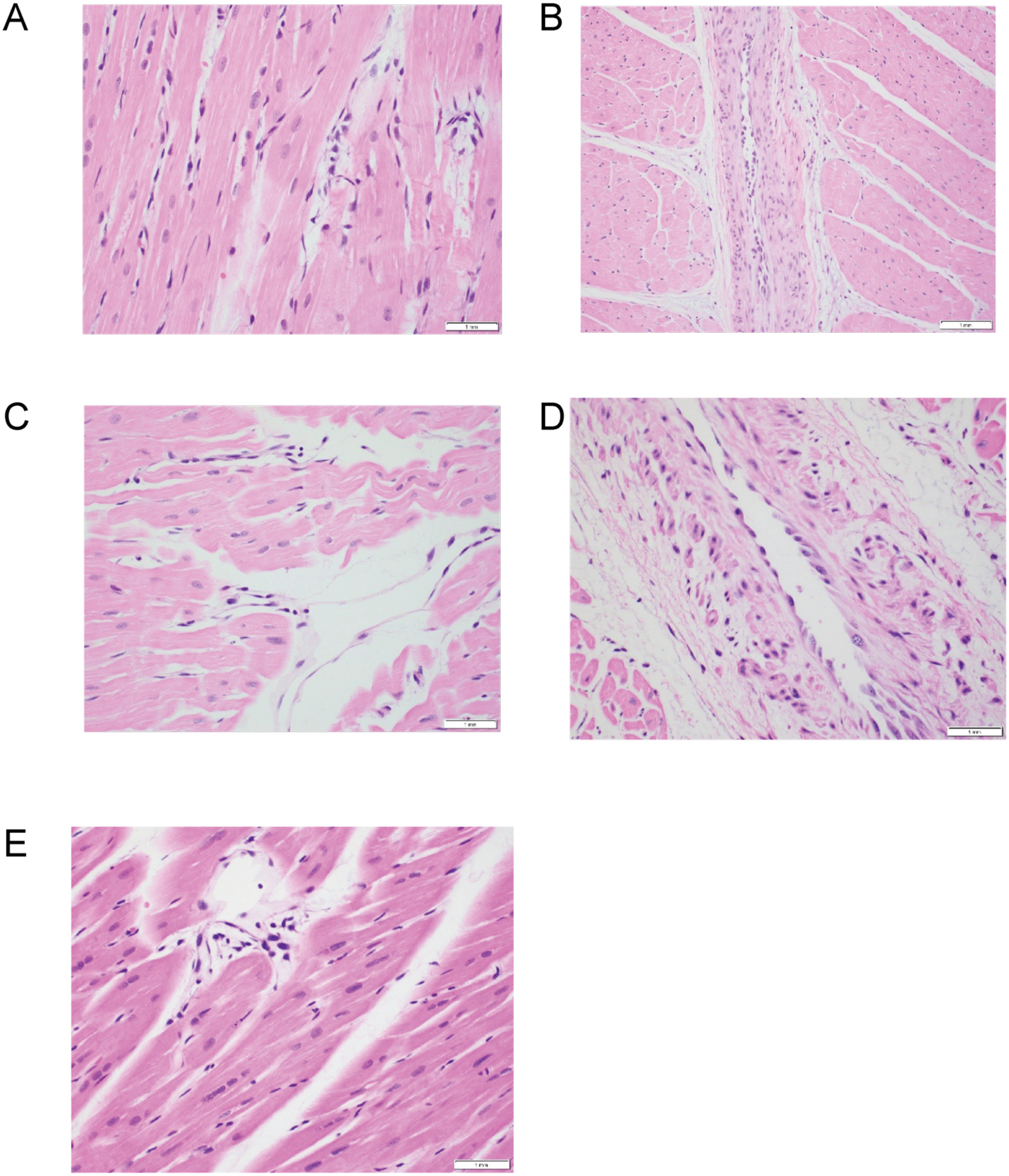


**Supplementary Figure 11:** H&E staining of Pig Xenograft Endomyocardial Biopsies from Decedent 1 (A-C) and Decedent 2 (D-E). A. Coagulative necrosis, which may be indicative of interstitial edema and Endothelial swelling of small interstitial capillary. B. Endothelial swelling of intramyocardial muscular arteries and perivascular edema. C. ‘Lifting’ of detached endothelial layers with few inflammatory cells evident. D. Endothelial swelling in intramyocardial muscular arteries. E. Endothelial swelling of small interstitial capillary

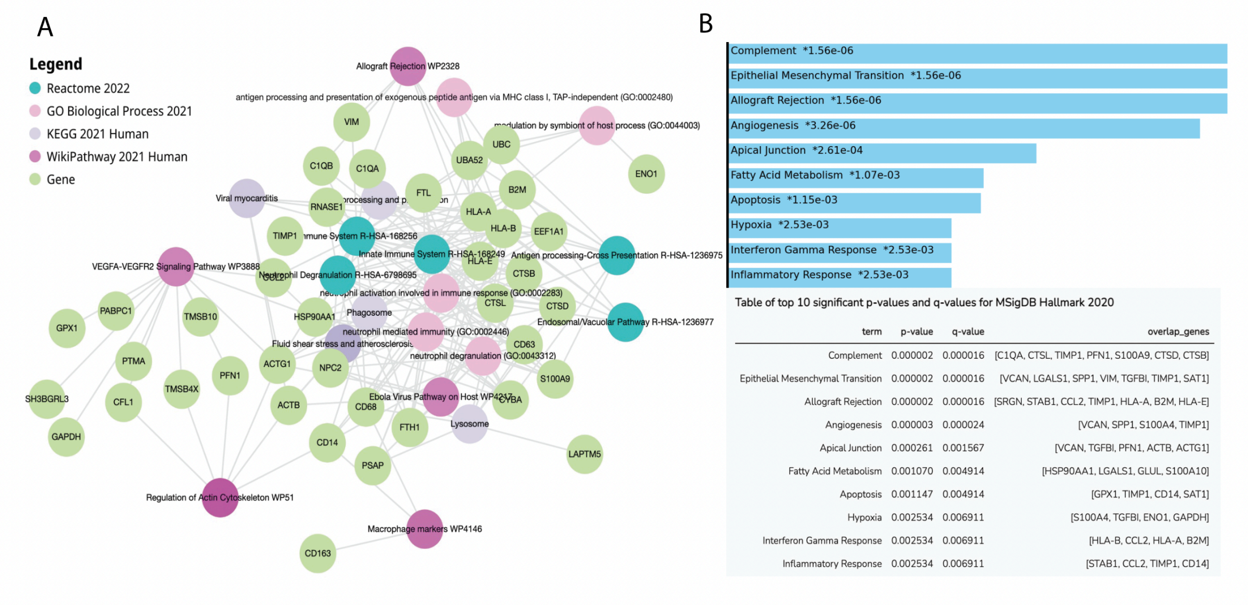


Table of top 10 significant p-values and q-values for MSigDB Hallmark 2020

| term | p-value | q-value | overlap_genes |
|---|---|---|---|
| Complement | 0.000002 | 0.000016 | [C1QA, CTSL, TIMP1, PFN1, S100A9, CTSD, CTSB] |
| Epithelial Mesenchymal Transition | 0.000002 | 0.000016 | [VCAN, LGALS1, SPP1, VIM, TGFBI, TIMP1, SAT1] |
| Allograft Rejection | 0.000002 | 0.000016 | [SRGN, STAB1, CCL2, TIMP1, HLA-A, B2M, HLA-E] |
| Angiogenesis | 0.000003 | 0.000024 | [VCAN, SPP1, S100A4, TIMP1] |
| Apical Junction | 0.000261 | 0.001567 | [VCAN, TGFBI, PFN1, ACTB, ACTG1] |
| Fatty Acid Metabolism | 0.001070 | 0.004914 | [HSP90AA1, LGALS1, GLUL, S100A10] |
| Apoptosis | 0.001147 | 0.004914 | [GPX1, TIMP1, CD14, SAT1] |
| Hypoxia | 0.002534 | 0.006911 | [S100A4, TGFBI, ENO1, GAPDH] |
| Interferon Gamma Response | 0.002534 | 0.006911 | [HLA-B, CCL2, HLA-A, B2M] |
| Inflammatory Response | 0.002534 | 0.006911 | [STAB1, CCL2, TIMP1, CD14] |

**Supplementary Figure 12.** Gene set enrichment analysis of the Visium cluster 6 (human cluster) marker genes (excluding mitochondrial, ribosomal genes and pig genes). The enrichment results, performed by enrichr-KG on 4 pathways gene set libraries, is shown in the format of a pathway-gene graph, with the top 5 gene set shown for each library. B. Enrichr gene-set enrichment analysis result of the same genes (Visium cluster 6 markers) in the MSigDB Hallmarks library, with a bar plot of the top enrichment and detailed table.

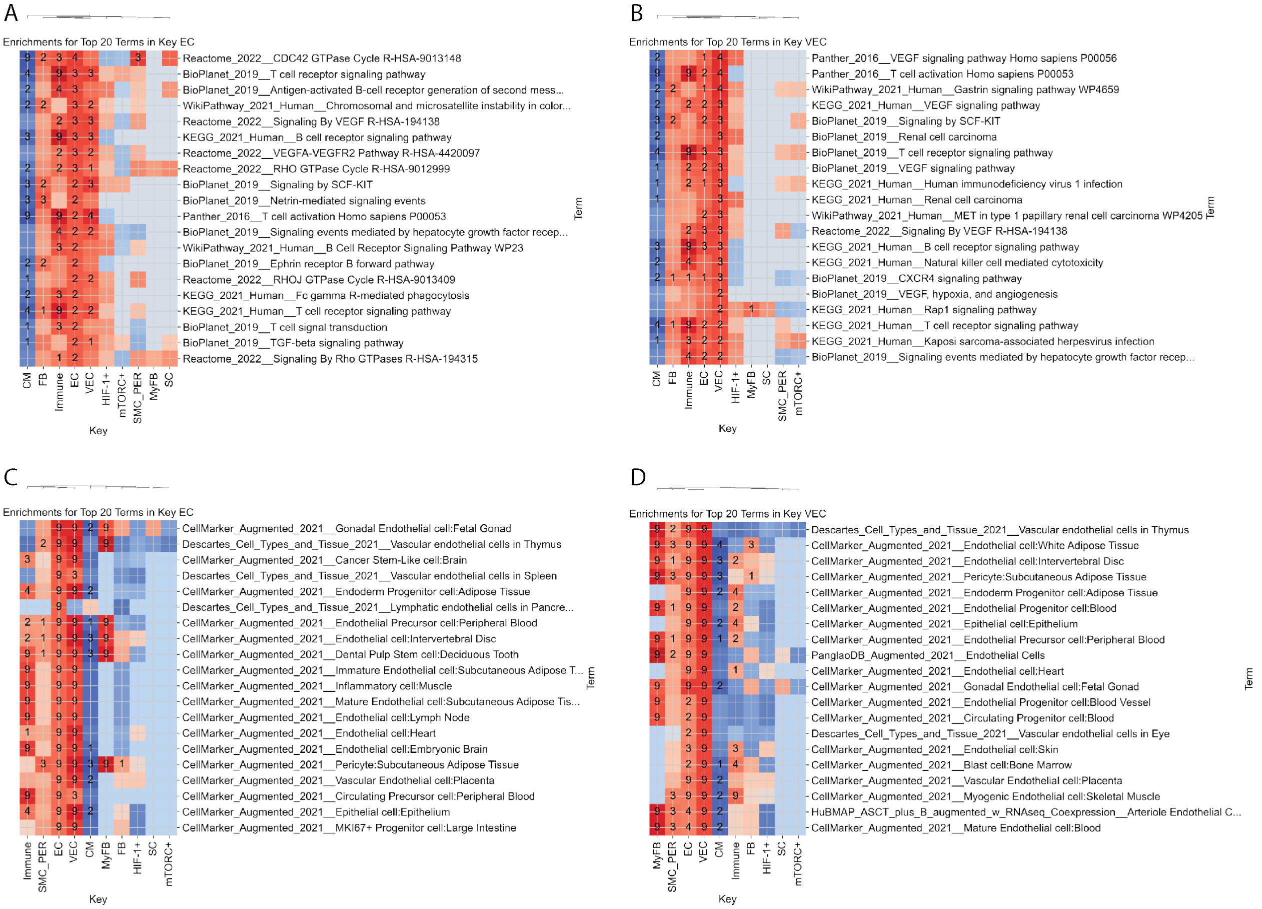


**Supplementary Figure 13.** Top 20 GSEA enrichment from Endothelial (EC, A and C) and Vascular Endothelial (VEC, B and D) cells, in the heart Xenograft tissue (pig cell), for pathways gene-sets (A and B) and cell-type gene-sets (C and D).

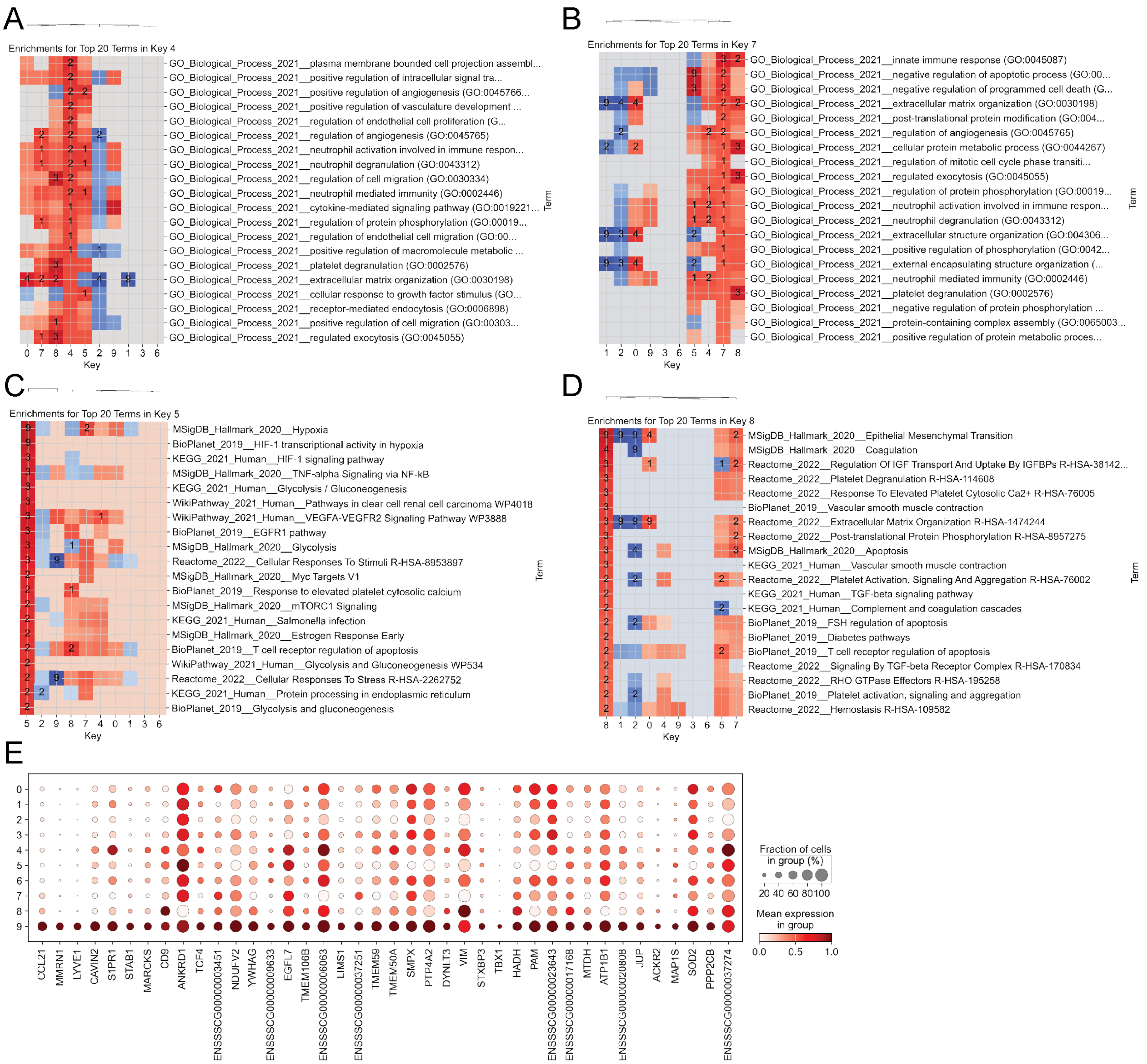


**Supplementary figure 14:** Understanding expression trends in Visium clusters. GO gene-set enrichment of cluster 4 (A) and 7 (B) markers. Pathway gene-set enrichment of cluster 5 (C) and 8 (D) markers. Main marker genes of cluster 9.

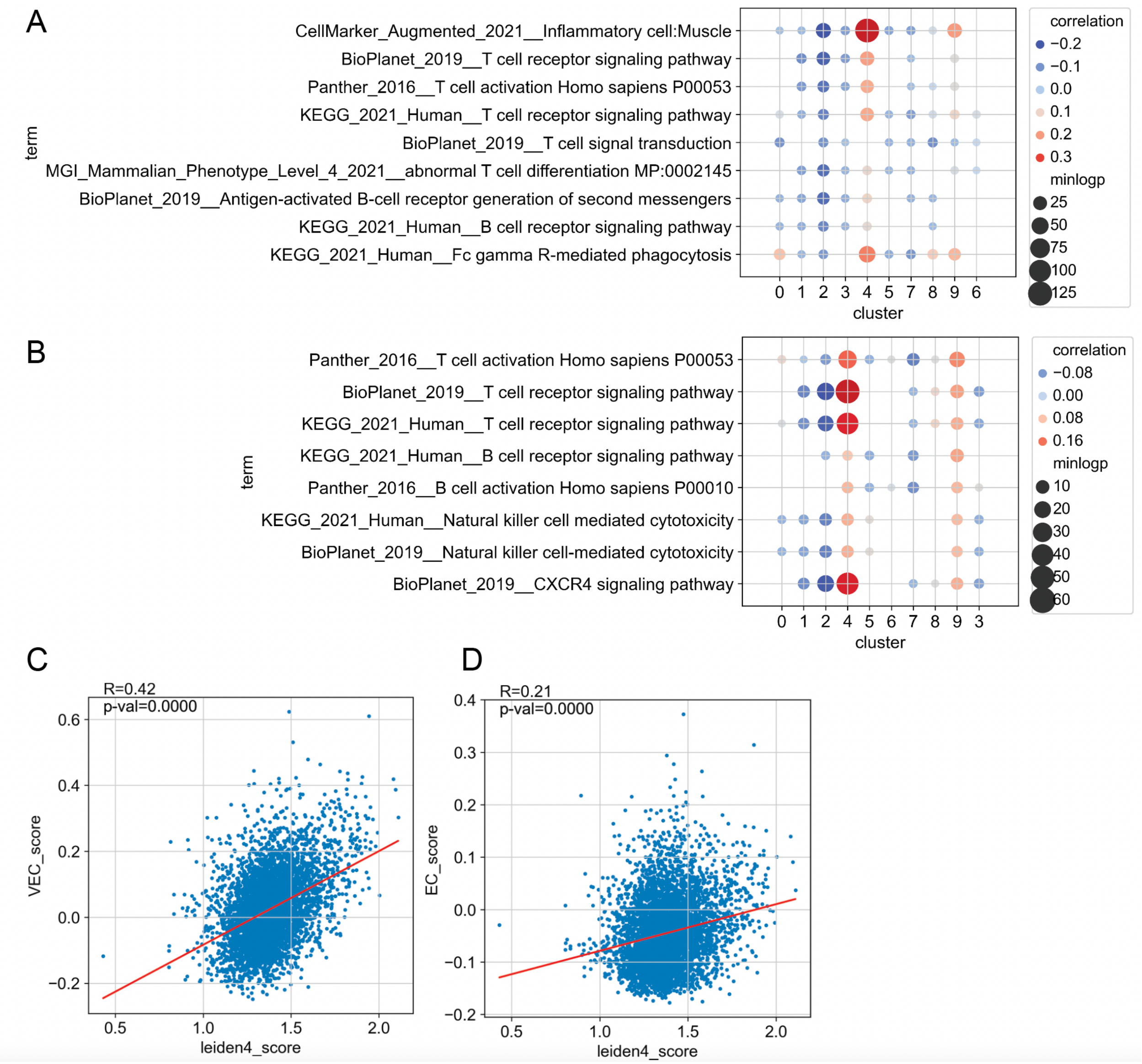


**Supplementary figure 15:** Immune related pathways found in EC and VEC (snRNA-seq) are found in the EC/VEC visium cluster 4. The correlation was performed between pathway score of Visium capture areas and score of each Visium group, within that group. The score is calculated independently in capture areas of each Visium cluster, for the pathways/lead genes from the EC immune enrichments (A) and VEC (B). Correlation between VEC (A) and EC (D) marker gene score within Visium capture areas, and the cluster 4 score.

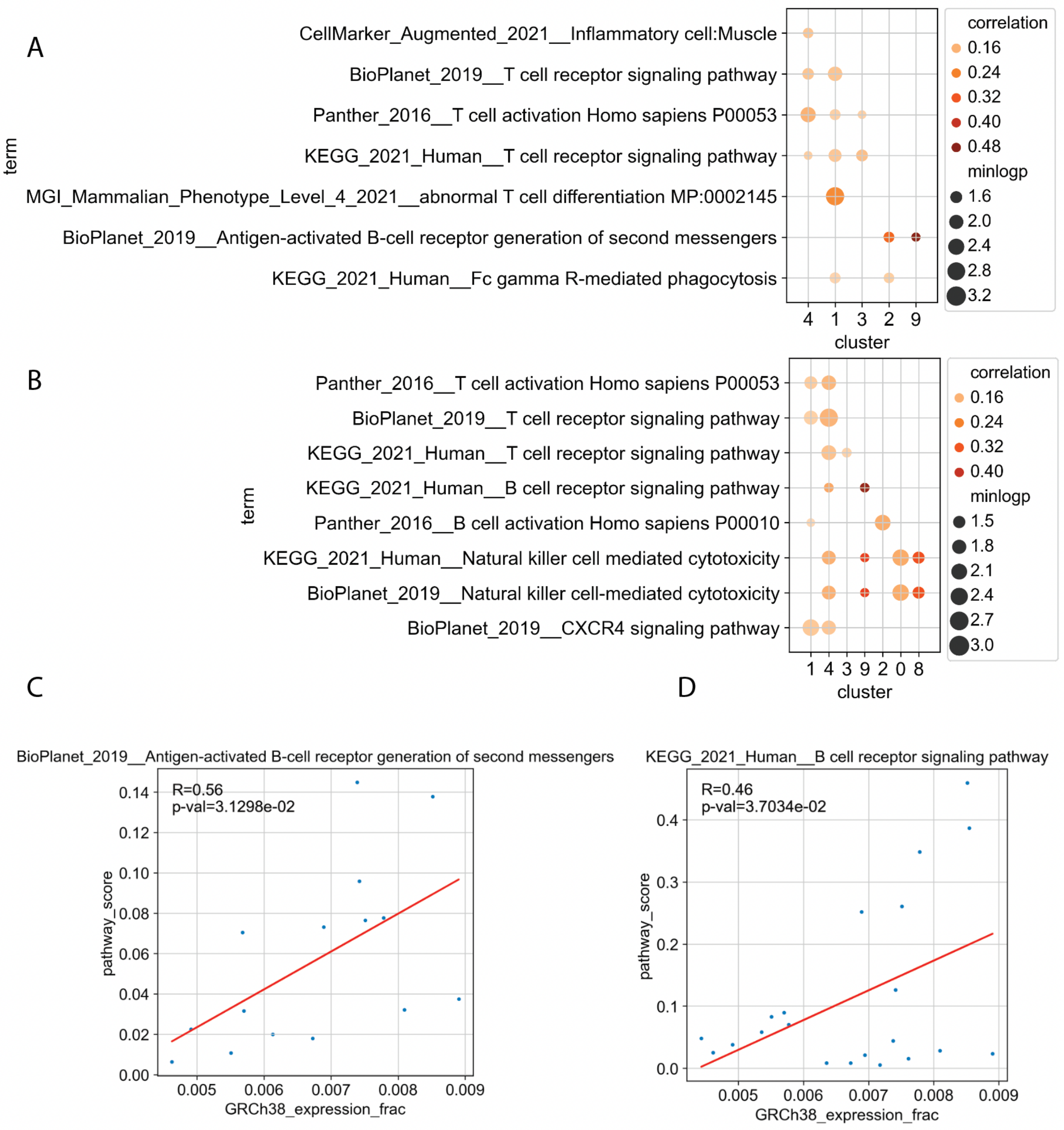


**Supplementary figure 16:** Correlation analysis of lead genes from immune related enrichments of Xenoheart endothelial populations (ECs and VECs). The correlation was performed between pathway score of Visium capture areas and fraction of counts aligning to the human genome. The score is calculated independently in capture areas of each Visium cluster, for the pathways/lead genes from the EC immune enrichments (A) and VEC (B). Only significant correlations (p-value < 0.05) are shown. Example of correlation within capture areas of the immune cluster (9), for Antigen-activated B-cell receptor generation of second messengers (C) and B cell receptor signaling pathway (D).
